# CCTP-Atlas: Compartment-Resolved Ciliary Transcriptional Dysregulation in Tetralogy of Fallot — A Six-Dataset Integrative Meta-Analysis

**DOI:** 10.64898/2026.10.12.758603

**Authors:** Rimsha Tahir, Talha Javed, Muhammad Moazzam Bin Rasheed, Qasim Aslam, Nadia Afsheen

**Author notes:** Corresponding author: Rimsha Tahir. Senior author: Nadia Afsheen.

## Abstract

Primary cilia coordinate Hedgehog-dependent outflow tract morphogenesis, and recent human genetics implicates ciliary loss-of-function in nonsyndromic tetralogy of Fallot (TOF). Yet compartment-resolved ciliary transcriptional programmes in public TOF single-nucleus data remain unmapped at module resolution. Here we curate a Cardiac Ciliary Transcriptomic Panel (CCTP; 1,611 genes; four modules) and execute a ten-phase integrative pipeline on six public sc/snRNA-seq resources. We harmonise published differential expression tables, tier evidence by recurrence across datasets, quantify compartment dysregulation with a permutation-validated Compartment Dysregulation Index (CMDI), assess cross-species directional concordance (CSCS), and align module activity with diffusion pseudotime in chambered cardiac organoids. Among 1,263 unique CCTP-associated hits, 396 genes recur in ≥2 dataset contrasts. A 24-gene TOF endothelial anchor includes PTPRM and FGFR1; only endothelial M1 structural-assembly CMDI exceeds a module-size-matched null (permutation p = 0.015). Patient-level pseudobulk (three TOF versus five donors) returns zero FDR-significant CCTP genes. CCTP-conditioned overlap with recent ciliary genetics is non-significant (Fisher p ≈ 0.13). Organoid pseudotime shows peak M1/M2 module scores during mid-maturation (culture day 8). CCTP-Atlas is a reproducible, tiered prioritisation resource; findings are hypothesis-generating and require expanded cohorts and functional validation.

## Introduction

Tetralogy of Fallot (TOF) is the most common cyanotic congenital heart defect. Surgical repair improves survival, yet long-term pulmonary, arrhythmia, and right ventricular complications remain frequent. Primary cilia transduce Hedgehog and related cues required for outflow tract development, ciliopathies often include congenital heart disease, and ciliary gene networks intersect endothelial–mesenchymal programmes relevant to RVOT maldevelopment (Goetz & Anderson, 2010; Djenoune et al., 2022). Concurrent exome evidence links ciliary loss-of-function variants to nonsyndromic TOF risk (Zhou et al., 2025), motivating compartment-resolved maps of ciliary transcription in human TOF tissue.

Single-nucleus RNA sequencing has mapped CHD cell states at scale (Hill et al., 2022; Zulibiya et al., 2023). Cardiac organoid models and cross-species atlases provide complementary developmental and regeneration contexts (Ho et al., 2022; Zhang et al., 2022; de Soysa et al., 2019; Li et al., 2025) but have not been integrated through a cilia-centric, module-resolved framework in TOF. Here we present CCTP-Atlas, an integrative computational study that (i) defines a reproducible CCTP reference and cross-dataset dysregulation catalogue, (ii) localises compartment-specific module dysregulation with CMDI, and (iii) relates human TOF signals to organoid maturation and partial cross-species concordance. Analyses are meta-analytic harmonisations of published gene lists rather than unified reprocessing of all raw count matrices.

## Results

### Integrative design and CCTP hit landscape

Six public datasets spanning TOF tissue, cardiac organoids, and cross-species references were integrated (Figure 1; Table 1). Across datasets, 1,263 unique genes intersected CCTP membership with dataset-specific dysregulation calls; 396 met a ≥2-dataset recurrence criterion (70 in ≥3). Hill et al. contributed the largest single-dataset footprint (1,229 unique CCTP hits); Ho organoids contributed 47 CCTP hits.

**Table 1.**
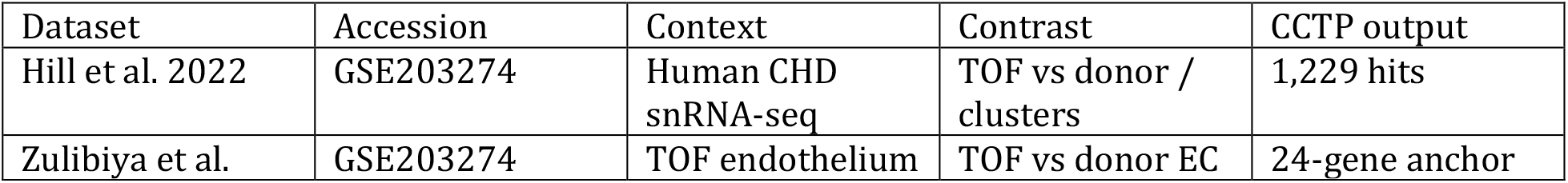
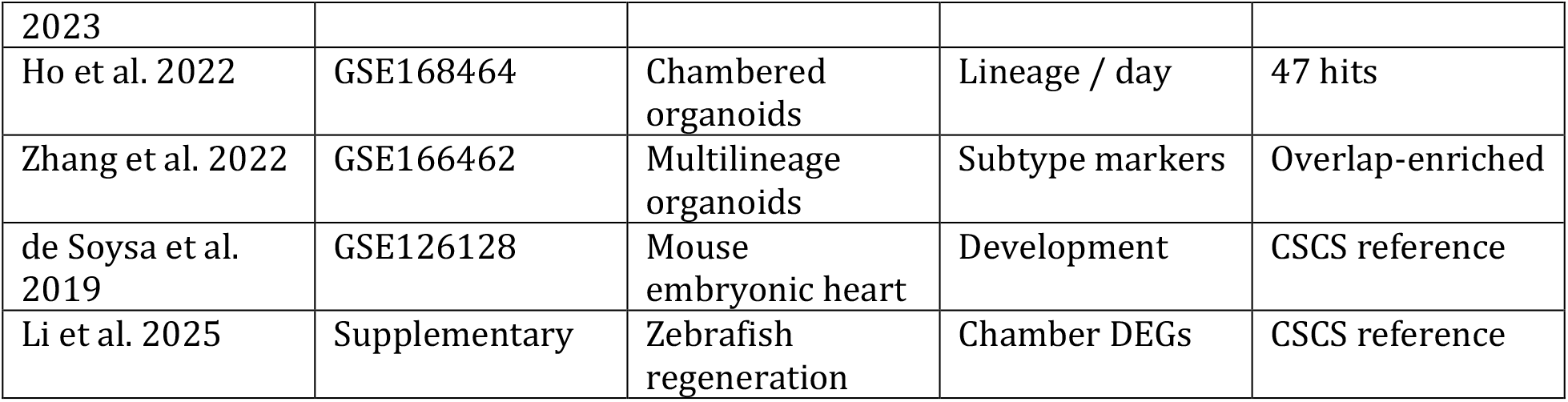
Integrated datasets.

**Figure 1.**
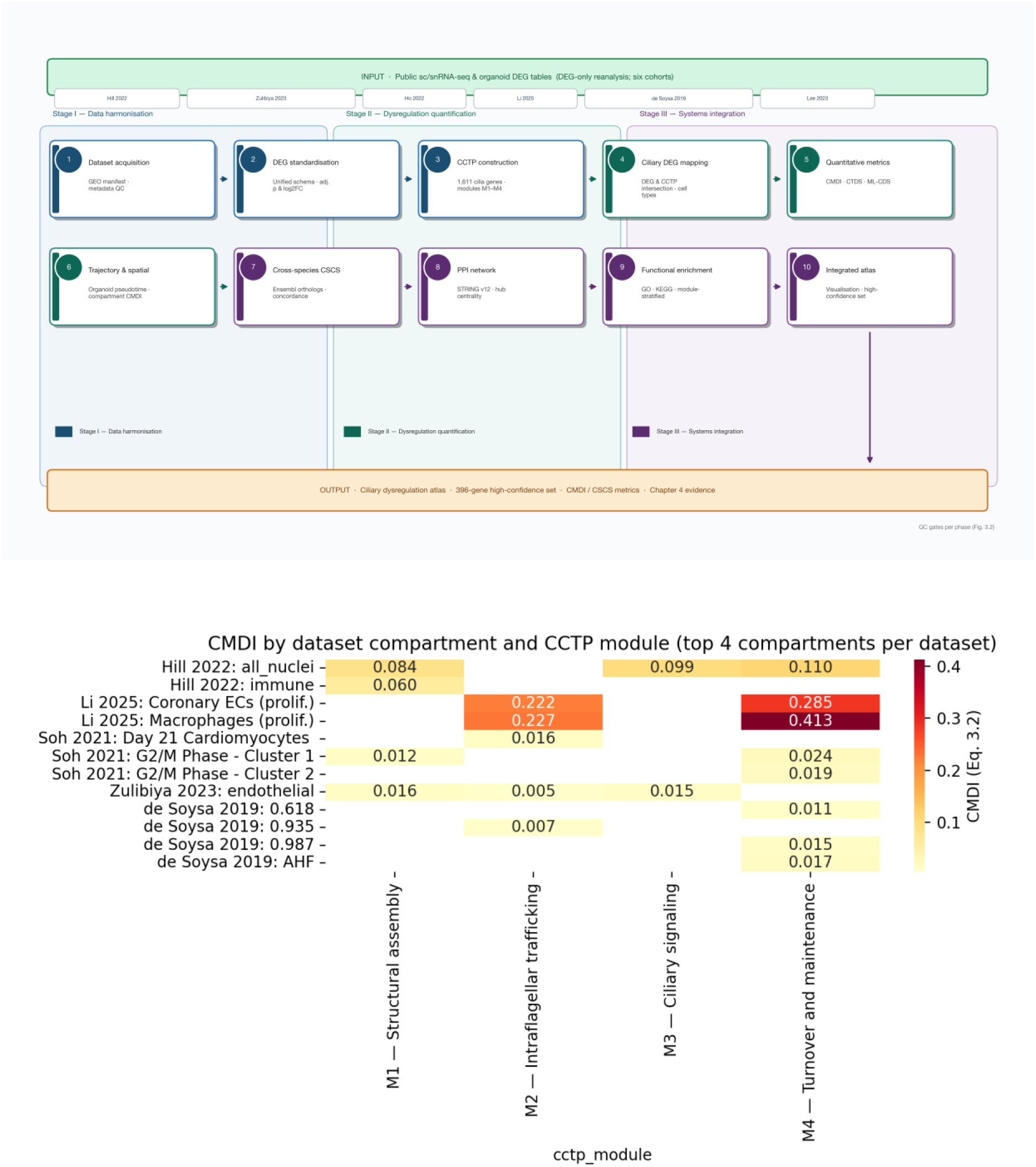
CCTP-Atlas overview. (A) Ten-phase integrative analysis pipeline. (B) CMDI across top compartment–module combinations; strongest endothelial M1 signal in Zulibiya TOF endothelium.

### TOF endothelial anchor and compartment-specific CMDI

Zulibiya endothelial TOF versus donor analysis yielded 24 genes passing adjusted p < 0.05 and |log2FC| ≥ 0.5 (Figure 2). PTPRM (M3; log2FC 2.61) and receptor tyrosine kinases FGFR1 and EGFR support endothelial RTK dysregulation; M1 structural genes (MAP4, ABLIM1, ARL15) indicate cytoskeletal and ciliogenesis remodelling. CMDI across 249 compartment–module combinations identified the strongest empirical signal for M1 in Zulibiya TOF endothelium (permutation p = 0.015; Figure 1B).

**Figure 2.**
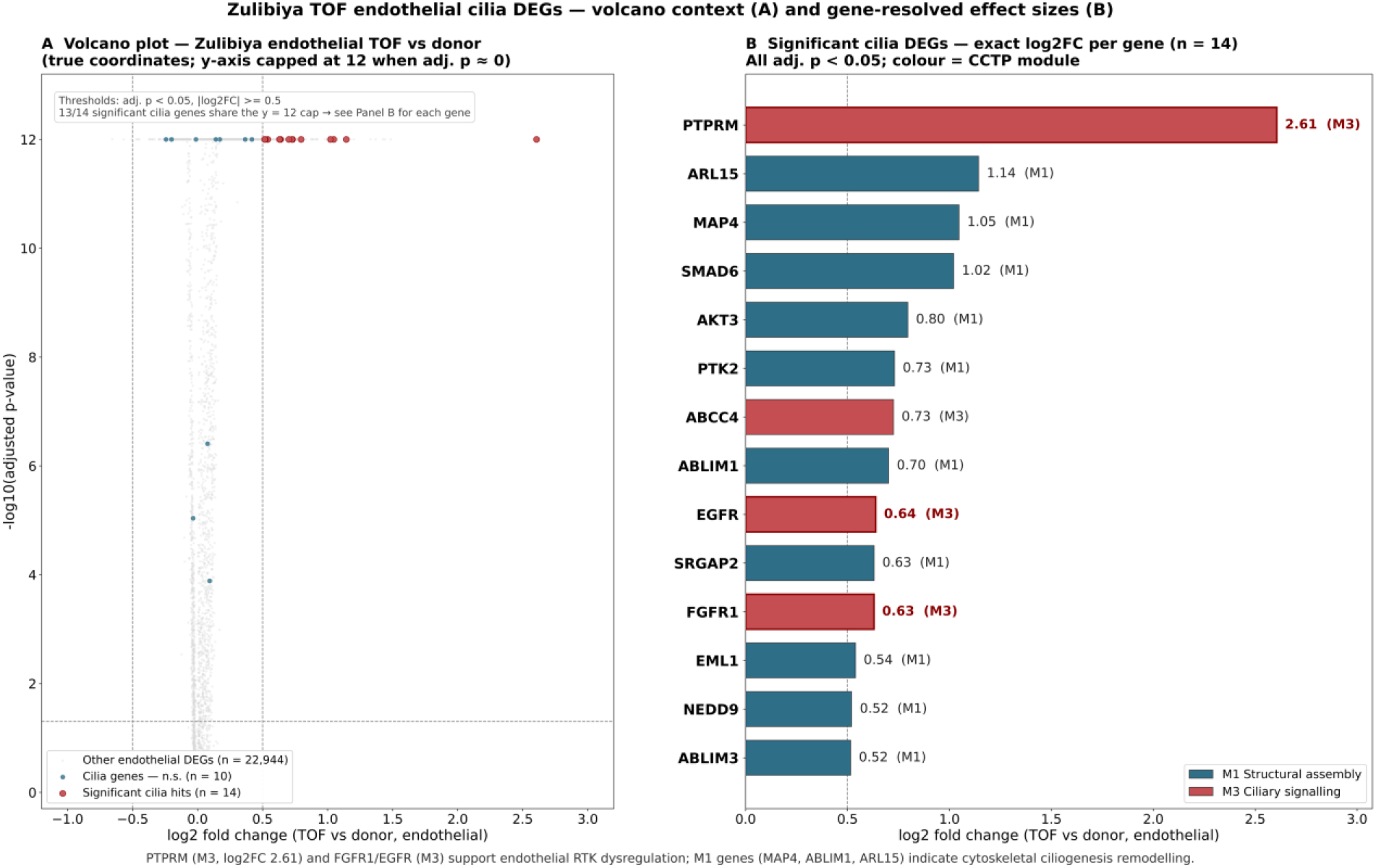
Zulibiya TOF endothelial cilia dysregulation. (A) Volcano plot (TOF vs donor endothelium). (B) Exact log2 fold changes for significant cilia DEGs coloured by CCTP module.

### Cross-species concordance, organoid pseudotime, and network architecture

Directional concordance among anchor genes was incomplete but informative (Figure 3A). Genome-wide CSCS across 396 high-confidence genes classified 11 as fully conserved and 136 as partially conserved (mean CSCS 0.69). On 17,290 Ho organoid cells, M1–M4 module scores peaked during mid-pseudotime (Figure 3B). STRING analysis (confidence ≥ 0.4) linked anchor genes to hubs including CDK1, EGFR, FGFR1, NOTCH1, and PTK2 (Figure 4). Patient-level pseudobulk returned zero FDR-significant CCTP genes; CCTP-conditioned overlap with Zhou et al. ciliary genetics was non-significant (Fisher p ≈ 0.13).

**Figure 3.**
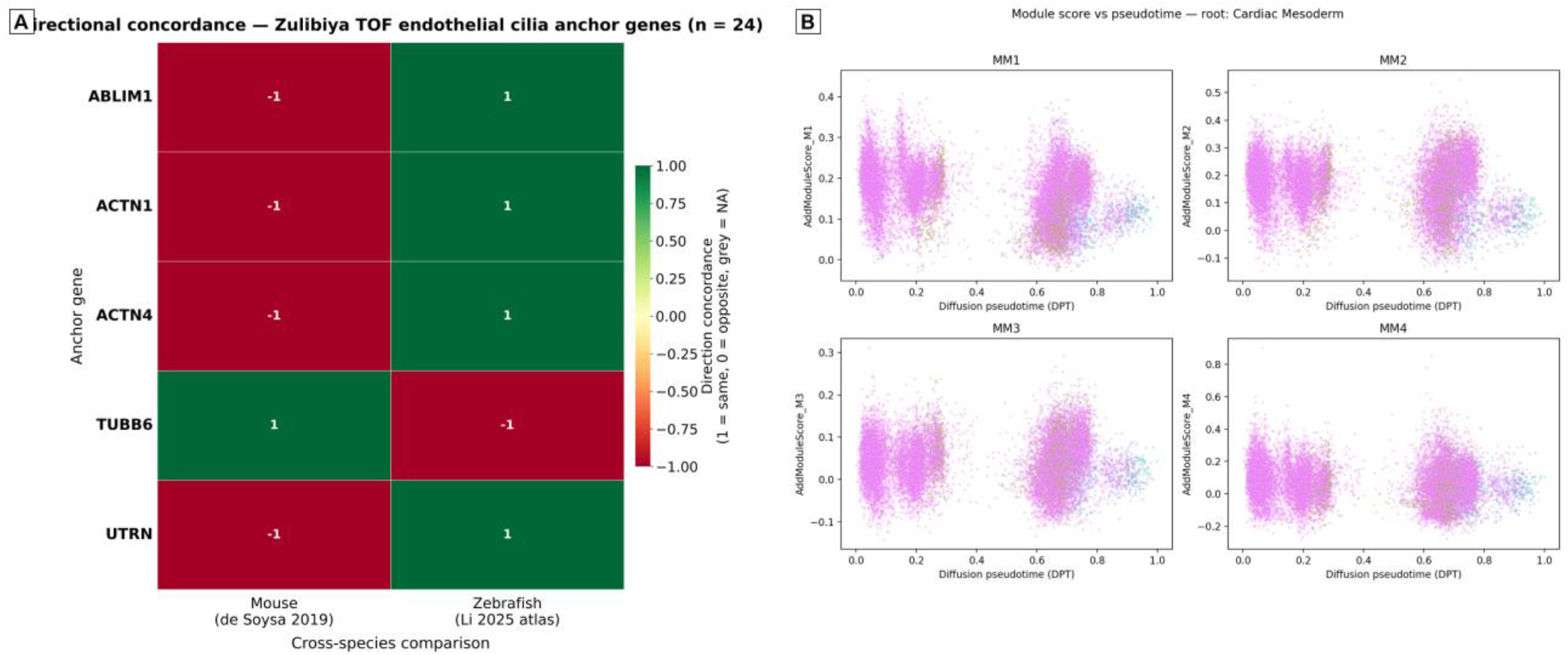
Cross-species and organoid context. (A) Directional concordance for anchor genes in mouse and zebrafish references. (B) CCTP module scores versus diffusion pseudotime in Ho chambered organoids.

**Figure 4.**
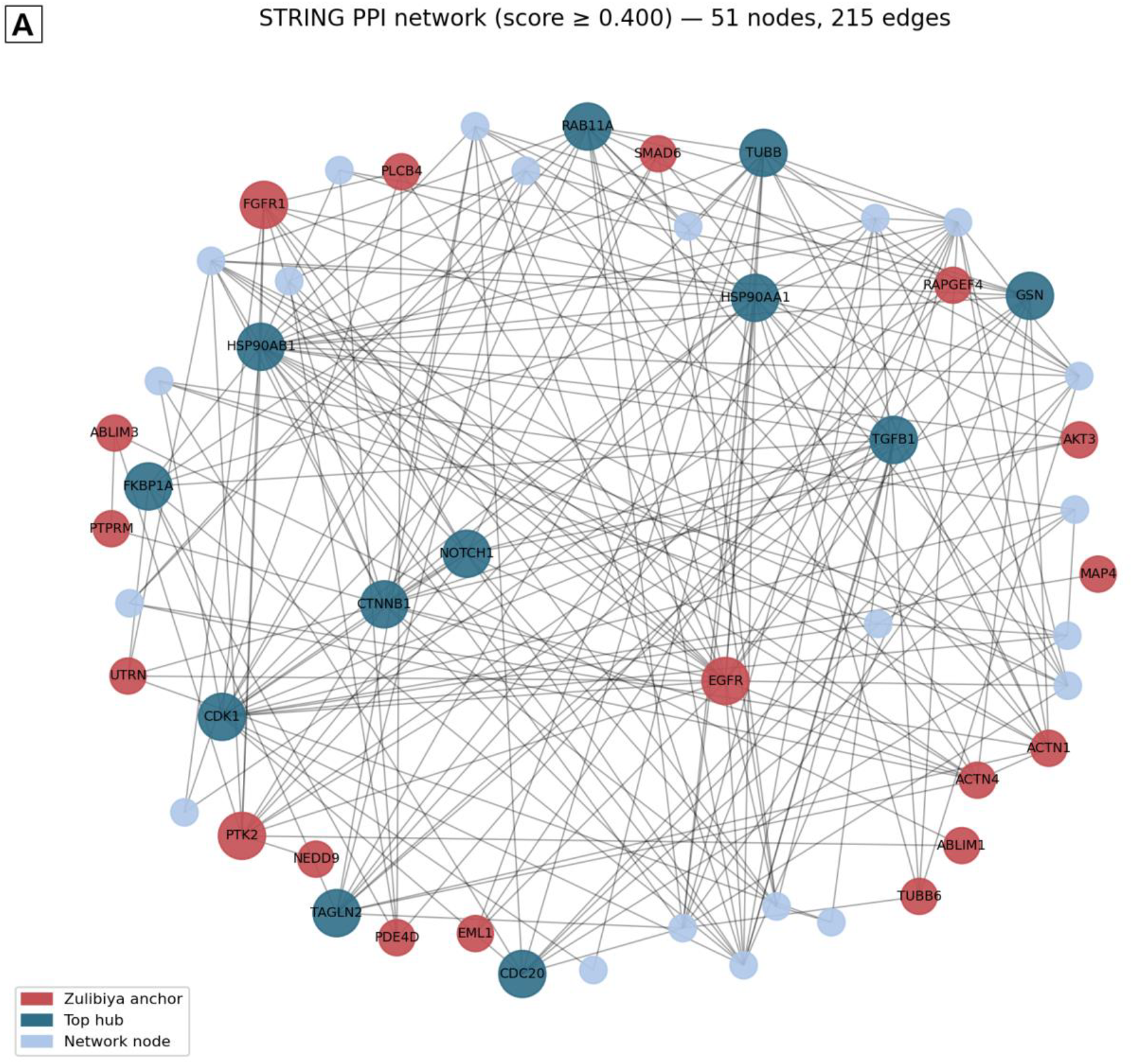
STRING protein–protein interaction network of high-confidence CCTP genes (confidence ≥ 0.4). Red nodes: Zulibiya anchor genes; teal nodes: top hubs.

## Discussion

CCTP-Atlas maps module-structured ciliary transcriptional correlates in TOF across human endothelial disease tissue, organoid maturation, and cross-species references. Dysregulation is compartment-specific, with peak CMDI in TOF endothelium rather than uniformly across cardiomyocyte clusters. Organoid pseudotime partially mirrors human module programmes but does not recapitulate disease endothelium. Cross-species concordance is selective, supporting prioritisation of cytoskeletal and adhesion nodes while leaving many TOF signals human-restricted.

Limitations include meta-analysis of published differential expression tables, partial independence because Hill and Zulibiya share GSE203274, marker-only organoid tables, and absence of experimental validation. Findings are hypothesis-generating. Priority follow-up includes endothelial PTPRM/FGFR1 localisation in TOF RVOT tissue, cilia metrics versus M1 scores in organoids, and perturbation in iPSC-derived endothelium.

## Materials and Methods

### Study design and datasets

Six public sc/snRNA-seq resources were integrated (Table 1). Hill et al. (2022) and Zulibiya et al. (2023) share GSE203274 but address distinct contrasts (multi-compartment CHD nuclei versus TOF versus donor endothelium). Ho et al. (2022; GSE168464) provided chambered organoid expression; Zhang et al. (2022; GSE166462) provided multilineage organoid markers; de Soysa et al. (2019; GSE126128) provided mouse developmental contrasts; Li et al. (2025) provided zebrafish chamber and regeneration FindMarkers. Drakhlis (GSE150017) and Hofbauer Visium cardioid spatial matrices were excluded when deposited accessions lacked analysable expression matrices (Supplementary Note S1).

### CCTP curation

The Cardiac Ciliary Transcriptomic Panel comprised 1,611 genes from primary cilia ontologies, intraflagellar transport components, ciliopathy gene lists, and cardiac developmental regulators. Genes were assigned to four modules: M1 structural assembly; M2 cell-cycle/proliferation; M3 ciliary signalling; M4 transport/IFT (Supplementary Table S1).

### DEG harmonisation and atlas construction

Published differential expression tables were standardised (gene symbol, log2FC, adjusted p-value, compartment). Thresholds: adjusted p < 0.05 and |log2FC| ≥ 0.5 where statistics were available. Marker-only tables without p-values were retained for overlap counting but excluded from volcano-based inference. High-confidence atlas genes required recurrence in ≥2 independent dataset contrasts. The 24-gene endothelial anchor was defined from Zulibiya TOF versus donor endothelium passing FDR and fold-change filters.

### Cross-Species Concordance Score (CSCS)

Ensembl REST API mapped human symbols to mouse and zebrafish orthologs. CSCS quantified directional concordance of dysregulation across species using harmonised DEG signs (Supplementary Methods).

### Compartment Dysregulation Index (CMDI)

CMDI summarised module-level dysregulation density per cell compartment using DEG-supported surrogates where per-cell matrices were unavailable. Empirical significance was assessed by label-permutation null models (10,000 iterations; fixed random seed).

### Organoid pseudotime

Ho merged UMI matrix (GSE168464) was processed in Scanpy (normalisation, HVG selection, PCA, neighbours, UMAP). Diffusion pseudotime was rooted at cardiac mesoderm clusters. Module scores were computed by sc.tl.score_genes for M1–M4 sets.

### Patient-level pseudobulk and genetics overlap

Where donor-level counts were available, pseudobulk aggregation compared three TOF versus five control donors with FDR control. Overlap between CCTP high-confidence genes and ciliary genetics loci (Zhou et al., 2025) was tested by Fisher’s exact test.

### Network and enrichment analysis

STRING v12.0 (confidence ≥ 0.4) built PPI networks for high-confidence genes. GO and KEGG enrichment used clusterProfiler with Benjamini–Hochberg FDR < 0.05.

### Reproducibility and ethics

Scripts, manifests (logs/*.json), and processed CSVs are version-controlled. This study used publicly available, de-identified data; no new human samples were collected.

## Data availability

Processed gene lists, manifests, and analysis scripts will be deposited on Zenodo.

Source datasets: GSE203274, GSE168464, GSE166462, GSE126128; Li et al. (2025) supplementary tables.

## Acknowledgments

We thank public dataset authors for open deposition of CHD and organoid atlases.

## Author contributions

R.T.: conceptualisation, data curation, formal analysis, visualisation, writing—original draft. T.J.: pipeline development, writing—review and editing. M.M.B.R.: bioinformatics and visualisation support, writing—review and editing. Q.A. and N.A.: supervision, methodology review, writing—review and editing.

