## Supplemental Data 1 for "CCTP-Atlas: Compartment-Resolved Ciliary Transcriptional Dysregulation in Tetralogy of Fallot — A Six-Dataset Integrative Meta-Analysis"

**Table S1. CCTP module gene sets**

| **CCTP module** | **N genes** | **Enrichment eligible** | **Gene preview (first 15)** |
| --- | --- | --- | --- |
| M1 — Structural assembly | 249 | True | MYH10; TUBB4B; ACTN1; ACTN2; AK1; ANXA1; BMP2; CDK1; CENPF; EZR; FSTL1; HMGB2; MARCKS; TAGLN2; TUBA1A; … (249 genes total) |
| M2 — Intraflagellar trafficking | 48 | True | ATP2A2; FBN2; CDC42EP5; DYNLL1; HSP90AB1; PHGDH; TPD52; ARL3; ASAP1; ATP2B4; ATP6V0D1; ATP6V1C1; ATP6V1D; COPS8; DAPK3; … (48 genes total) |
| M3 — Ciliary signaling | 72 | True | PKIG; TGFB1; ABCC4; CTNNB1; EGFR; FGFR1; NOTCH1; PDGFRA; TEK; ACVR1B; ADAM17; ADCY6; ADORA2A; ANO10; ARHGAP35; … (72 genes total) |
| M4 — Turnover and maintenance | 21 | True | RAB11A; PLK1; PSMC2; PSMC5; ATG16L1; ATG3; ATG5; AURKA; CHMP4B; HUWE1; MCM2; NEK2; PSMB7; RAB7A; SIAH2; … (21 genes total) |
| M7 — Motile cilia (non-primary) | 6 | True | CFAP70; DYNC1I2; IFT140; IFT57; IFT74; SPAG1 |

**Table S2. 396 high-confidence atlas genes (recurrent in ≥2 datasets)**

| **Gene** | **N datasets** | **Datasets** | **CCTP module** | **Cilia module** | **In 24-gene anchor** |
| --- | --- | --- | --- | --- | --- |
| ATP2A2 | 5 | de_soysa_2019;hill_2022;lee_2023;li_2025;soh_2021 | M2 — Intraflagellar trafficking | 3 IFT/Trafficking/BBSome | False |
| MYH10 | 5 | de_soysa_2019;hill_2022;lee_2023;li_2025;soh_2021 | M1 — Structural assembly | 6 Cytoskeleton ciliogenesis regulation | False |
| TUBB4B | 5 | de_soysa_2019;hill_2022;lee_2023;li_2025;soh_2021 | M1 — Structural assembly | 1 Structural | False |
| ACTN1 | 4 | hill_2022;lee_2023;li_2025;zulibiya_2023 | M1 — Structural assembly | 6 Cytoskeleton ciliogenesis regulation | True |
| ACTN2 | 4 | de_soysa_2019;hill_2022;li_2025;soh_2021 | M1 — Structural assembly | 6 Cytoskeleton ciliogenesis regulation | False |
| AK1 | 4 | de_soysa_2019;hill_2022;li_2025;soh_2021 | M1 — Structural assembly | 1 Structural | False |
| ANXA1 | 4 | hill_2022;lee_2023;li_2025;soh_2021 | M1 — Structural assembly | 1 Structural | False |
| BMP2 | 4 | de_soysa_2019;hill_2022;lee_2023;soh_2021 | M1 — Structural assembly | 1 Structural | False |
| CDK1 | 4 | hill_2022;lee_2023;li_2025;soh_2021 | M1 — Structural assembly | 1 Structural | False |
| CENPF | 4 | hill_2022;lee_2023;li_2025;soh_2021 | M1 — Structural assembly | 1 Structural | False |
| EZR | 4 | de_soysa_2019;hill_2022;li_2025;soh_2021 | M1 — Structural assembly | 6 Cytoskeleton ciliogenesis regulation | False |
| FBN2 | 4 | de_soysa_2019;hill_2022;lee_2023;soh_2021 | M2 — Intraflagellar trafficking | 3 IFT/Trafficking/BBSome | False |
| FSTL1 | 4 | de_soysa_2019;hill_2022;lee_2023;li_2025 | M1 — Structural assembly | 2 Assembly | False |
| HMGB2 | 4 | hill_2022;lee_2023;li_2025;soh_2021 | M1 — Structural assembly | 1 Structural | False |
| MARCKS | 4 | de_soysa_2019;hill_2022;lee_2023;soh_2021 | M1 — Structural assembly | 6 Cytoskeleton ciliogenesis regulation | False |
| PKIG | 4 | hill_2022;lee_2023;li_2025;soh_2021 | M3 — Ciliary signaling | 4 Signaling Associated | False |
| RAB11A | 4 | de_soysa_2019;hill_2022;li_2025;soh_2021 | M4 — Turnover and maintenance | 5 Turnover/Maintenance | False |
| TAGLN2 | 4 | de_soysa_2019;hill_2022;lee_2023;soh_2021 | M1 — Structural assembly | 1 Structural | False |
| TGFB1 | 4 | de_soysa_2019;hill_2022;li_2025;soh_2021 | M3 — Ciliary signaling | 4 Signaling Associated | False |
| TUBA1A | 4 | de_soysa_2019;hill_2022;li_2025;soh_2021 | M1 — Structural assembly | 1 Structural | False |
| TUBA1B | 4 | hill_2022;lee_2023;li_2025;soh_2021 | M1 — Structural assembly | 1 Structural | False |
| TUBB | 4 | hill_2022;lee_2023;li_2025;soh_2021 | M1 — Structural assembly | 1 Structural | False |
| TUBB6 | 4 | de_soysa_2019;hill_2022;li_2025;zulibiya_2023 | M1 — Structural assembly | 6 Cytoskeleton ciliogenesis regulation | True |
| ABCC4 | 3 | hill_2022;li_2025;zulibiya_2023 | M3 — Ciliary signaling | 4 Signaling Associated | True |
| ABLIM1 | 3 | hill_2022;li_2025;zulibiya_2023 | M1 — Structural assembly | 6 Cytoskeleton ciliogenesis regulation | True |
| ABLIM3 | 3 | hill_2022;li_2025;zulibiya_2023 | M1 — Structural assembly | 6 Cytoskeleton ciliogenesis regulation | True |
| ACTN4 | 3 | hill_2022;li_2025;zulibiya_2023 | M1 — Structural assembly | 6 Cytoskeleton ciliogenesis regulation | True |
| ANXA6 | 3 | de_soysa_2019;hill_2022;li_2025 | M1 — Structural assembly | 1 Structural | False |
| APP | 3 | hill_2022;lee_2023;li_2025 | M1 — Structural assembly | 1 Structural | False |
| CDC20 | 3 | hill_2022;lee_2023;soh_2021 | M1 — Structural assembly | 1 Structural | False |
| CDC42EP5 | 3 | hill_2022;li_2025;soh_2021 | M2 — Intraflagellar trafficking | 3 IFT/Trafficking/BBSome | False |
| CEP55 | 3 | hill_2022;li_2025;soh_2021 | M1 — Structural assembly | 1 Structural | False |
| CKAP2 | 3 | hill_2022;li_2025;soh_2021 | M1 — Structural assembly | 6 Cytoskeleton ciliogenesis regulation | False |
| CTNNB1 | 3 | hill_2022;lee_2023;li_2025 | M3 — Ciliary signaling | 4 Signaling Associated | False |
| DYNLL1 | 3 | hill_2022;lee_2023;li_2025 | M2 — Intraflagellar trafficking | 3 IFT/Trafficking/BBSome | False |
| EEF1A1 | 3 | hill_2022;lee_2023;li_2025 | M1 — Structural assembly | 2 Assembly | False |
| EGFR | 3 | hill_2022;li_2025;zulibiya_2023 | M3 — Ciliary signaling | 4 Signaling Associated | True |
| EHD4 | 3 | de_soysa_2019;hill_2022;li_2025 | M1 — Structural assembly | 1 Structural | False |
| EML1 | 3 | hill_2022;li_2025;zulibiya_2023 | M1 — Structural assembly | 6 Cytoskeleton ciliogenesis regulation | True |
| FGFR1 | 3 | hill_2022;li_2025;zulibiya_2023 | M3 — Ciliary signaling | 4 Signaling Associated | True |
| FKBP1A | 3 | hill_2022;lee_2023;soh_2021 | M1 — Structural assembly | 1 Structural | False |
| GNAI1 | 3 | de_soysa_2019;hill_2022;li_2025 | M1 — Structural assembly | 1 Structural | False |
| GSN | 3 | de_soysa_2019;hill_2022;li_2025 | M1 — Structural assembly | 1 Structural | False |
| HK1 | 3 | de_soysa_2019;hill_2022;li_2025 | M1 — Structural assembly | 1 Structural | False |
| HSP90AA1 | 3 | hill_2022;li_2025;soh_2021 | M1 — Structural assembly | 2 Assembly | False |
| HSP90AB1 | 3 | hill_2022;li_2025;soh_2021 | M2 — Intraflagellar trafficking | 3 IFT/Trafficking/BBSome | False |
| KIF20B | 3 | hill_2022;li_2025;soh_2021 | M1 — Structural assembly | 1 Structural | False |
| LRRC49 | 3 | hill_2022;lee_2023;soh_2021 | M1 — Structural assembly | 1 Structural | False |
| NEDD9 | 3 | hill_2022;li_2025;zulibiya_2023 | M1 — Structural assembly | 1 Structural | True |
| NOTCH1 | 3 | de_soysa_2019;hill_2022;li_2025 | M3 — Ciliary signaling | 4 Signaling Associated | False |
| NUAK1 | 3 | hill_2022;li_2025;zulibiya_2023 | M1 — Structural assembly | 6 Cytoskeleton ciliogenesis regulation | True |
| PDGFRA | 3 | hill_2022;li_2025;soh_2021 | M3 — Ciliary signaling | 4 Signaling Associated | False |
| PHGDH | 3 | de_soysa_2019;hill_2022;soh_2021 | M2 — Intraflagellar trafficking | 3 IFT/Trafficking/BBSome | False |
| PITX2 | 3 | de_soysa_2019;hill_2022;lee_2023 | M1 — Structural assembly | 1 Structural | False |
| PLK1 | 3 | hill_2022;li_2025;soh_2021 | M4 — Turnover and maintenance | 5 Turnover/Maintenance | False |
| PRDX1 | 3 | hill_2022;li_2025;soh_2021 | M1 — Structural assembly | 2 Assembly | False |
| PSMC2 | 3 | de_soysa_2019;hill_2022;li_2025 | M4 — Turnover and maintenance | 5 Turnover/Maintenance | False |
| PSMC4 | 3 | de_soysa_2019;hill_2022;li_2025 | M1 — Structural assembly | 1 Structural | False |
| PSMC5 | 3 | de_soysa_2019;hill_2022;li_2025 | M4 — Turnover and maintenance | 5 Turnover/Maintenance | False |
| RCSD1 | 3 | de_soysa_2019;hill_2022;li_2025 | M1 — Structural assembly | 6 Cytoskeleton ciliogenesis regulation | False |
| RRAD | 3 | de_soysa_2019;hill_2022;li_2025 | M1 — Structural assembly | 1 Structural | False |
| SCCPDH | 3 | de_soysa_2019;hill_2022;li_2025 | M1 — Structural assembly | 1 Structural | False |
| SKP1 | 3 | hill_2022;lee_2023;li_2025 | M1 — Structural assembly | 1 Structural | False |
| SMAD6 | 3 | hill_2022;li_2025;zulibiya_2023 | M1 — Structural assembly | 1 Structural | True |
| TACC3 | 3 | hill_2022;li_2025;soh_2021 | M1 — Structural assembly | 6 Cytoskeleton ciliogenesis regulation | False |
| TEK | 3 | de_soysa_2019;hill_2022;li_2025 | M3 — Ciliary signaling | 4 Signaling Associated | False |
| TPD52 | 3 | de_soysa_2019;hill_2022;li_2025 | M2 — Intraflagellar trafficking | 3 IFT/Trafficking/BBSome | False |
| TPPP3 | 3 | hill_2022;li_2025;soh_2021 | M1 — Structural assembly | 6 Cytoskeleton ciliogenesis regulation | False |
| TUBA1C | 3 | hill_2022;li_2025;soh_2021 | M1 — Structural assembly | 1 Structural | False |
| UTRN | 3 | hill_2022;li_2025;zulibiya_2023 | M1 — Structural assembly | 1 Structural | True |
| AAMP | 2 | hill_2022;li_2025 | M1 — Structural assembly | 6 Cytoskeleton ciliogenesis regulation | False |
| ABCC3 | 2 | hill_2022;li_2025 | M1 — Structural assembly | 1 Structural | False |
| ACAA2 | 2 | hill_2022;li_2025 | M1 — Structural assembly | 1 Structural | False |
| ACLY | 2 | hill_2022;li_2025 | M1 — Structural assembly | 1 Structural | False |
| ACTR2 | 2 | hill_2022;li_2025 | M1 — Structural assembly | 6 Cytoskeleton ciliogenesis regulation | False |
| ACTR3 | 2 | hill_2022;li_2025 | M1 — Structural assembly | 6 Cytoskeleton ciliogenesis regulation | False |
| ACVR1B | 2 | hill_2022;li_2025 | M3 — Ciliary signaling | 4 Signaling Associated | False |
| ADAM17 | 2 | hill_2022;li_2025 | M3 — Ciliary signaling | 4 Signaling Associated | False |
| ADAMTS9 | 2 | hill_2022;zulibiya_2023 | M1 — Structural assembly | 2 Assembly | True |
| ADCY6 | 2 | hill_2022;li_2025 | M3 — Ciliary signaling | 4 Signaling Associated | False |
| ADORA2A | 2 | hill_2022;li_2025 | M3 — Ciliary signaling | 4 Signaling Associated | False |
| ADRB3 | 2 | hill_2022;li_2025 | M1 — Structural assembly | 1 Structural | False |
| AK2 | 2 | hill_2022;li_2025 | M1 — Structural assembly | 1 Structural | False |
| AKT3 | 2 | hill_2022;zulibiya_2023 | M1 — Structural assembly | 1 Structural | True |
| ALDH1A1 | 2 | hill_2022;soh_2021 | M1 — Structural assembly | 1 Structural | False |
| ANO10 | 2 | hill_2022;li_2025 | M3 — Ciliary signaling | 4 Signaling Associated | False |
| APOO | 2 | hill_2022;li_2025 | M1 — Structural assembly | 1 Structural | False |
| ARHGAP1 | 2 | hill_2022;li_2025 | M1 — Structural assembly | 6 Cytoskeleton ciliogenesis regulation | False |
| ARHGAP35 | 2 | hill_2022;li_2025 | M3 — Ciliary signaling | 4 Signaling Associated | False |
| ARL15 | 2 | hill_2022;zulibiya_2023 | M1 — Structural assembly | 1 Structural | True |
| ARL2 | 2 | hill_2022;li_2025 | M1 — Structural assembly | 1 Structural | False |
| ARL3 | 2 | hill_2022;li_2025 | M2 — Intraflagellar trafficking | 3 IFT/Trafficking/BBSome | False |
| ARPP19 | 2 | hill_2022;li_2025 | M1 — Structural assembly | 1 Structural | False |
| ASAP1 | 2 | hill_2022;li_2025 | M2 — Intraflagellar trafficking | 3 IFT/Trafficking/BBSome | False |
| ATF3 | 2 | hill_2022;li_2025 | M1 — Structural assembly | 1 Structural | False |
| ATF6 | 2 | hill_2022;li_2025 | M1 — Structural assembly | 6 Cytoskeleton ciliogenesis regulation | False |
| ATG16L1 | 2 | hill_2022;li_2025 | M4 — Turnover and maintenance | 5 Turnover/Maintenance | False |
| ATG3 | 2 | hill_2022;li_2025 | M4 — Turnover and maintenance | 5 Turnover/Maintenance | False |
| ATG5 | 2 | hill_2022;li_2025 | M4 — Turnover and maintenance | 5 Turnover/Maintenance | False |
| ATP2B4 | 2 | hill_2022;li_2025 | M2 — Intraflagellar trafficking | 3 IFT/Trafficking/BBSome | False |
| ATP6V0D1 | 2 | hill_2022;li_2025 | M2 — Intraflagellar trafficking | 3 IFT/Trafficking/BBSome | False |
| ATP6V1C1 | 2 | hill_2022;li_2025 | M2 — Intraflagellar trafficking | 3 IFT/Trafficking/BBSome | False |
| ATP6V1D | 2 | hill_2022;li_2025 | M2 — Intraflagellar trafficking | 3 IFT/Trafficking/BBSome | False |
| AURKA | 2 | li_2025;soh_2021 | M4 — Turnover and maintenance | 5 Turnover/Maintenance | False |
| BAG2 | 2 | hill_2022;li_2025 | M1 — Structural assembly | 1 Structural | False |
| BAG3 | 2 | hill_2022;li_2025 | M1 — Structural assembly | 1 Structural | False |
| BCL3 | 2 | hill_2022;li_2025 | M1 — Structural assembly | 1 Structural | False |
| BUB1B | 2 | hill_2022;li_2025 | M1 — Structural assembly | 1 Structural | False |
| C7ORF57 | 2 | hill_2022;li_2025 | M1 — Structural assembly | 1 Structural | False |
| CACNA1C | 2 | hill_2022;li_2025 | M3 — Ciliary signaling | 4 Signaling Associated | False |
| CAMK1D | 2 | hill_2022;li_2025 | M1 — Structural assembly | 1 Structural | False |
| CAMK2A | 2 | hill_2022;li_2025 | M1 — Structural assembly | 1 Structural | False |
| CAMK2B | 2 | hill_2022;li_2025 | M1 — Structural assembly | 1 Structural | False |
| CAMK2G | 2 | hill_2022;li_2025 | M1 — Structural assembly | 1 Structural | False |
| CAMK2N2 | 2 | hill_2022;li_2025 | M1 — Structural assembly | 1 Structural | False |
| CATIP | 2 | hill_2022;li_2025 | M3 — Ciliary signaling | 4 Signaling Associated | False |
| CAV1 | 2 | hill_2022;li_2025 | M3 — Ciliary signaling | 4 Signaling Associated | False |
| CCSAP | 2 | hill_2022;li_2025 | M1 — Structural assembly | 1 Structural | False |
| CCT8 | 2 | hill_2022;li_2025 | M1 — Structural assembly | 1 Structural | False |
| CDH23 | 2 | hill_2022;li_2025 | M3 — Ciliary signaling | 4 Signaling Associated | False |
| CDK2 | 2 | hill_2022;li_2025 | M1 — Structural assembly | 1 Structural | False |
| CDK5RAP2 | 2 | hill_2022;li_2025 | M1 — Structural assembly | 2 Assembly | False |
| CDKN1B | 2 | hill_2022;li_2025 | M1 — Structural assembly | 6 Cytoskeleton ciliogenesis regulation | False |
| CEP170 | 2 | hill_2022;li_2025 | M1 — Structural assembly | 6 Cytoskeleton ciliogenesis regulation | False |
| CEP350 | 2 | hill_2022;li_2025 | M1 — Structural assembly | 1 Structural | False |
| CEP70 | 2 | hill_2022;li_2025 | M1 — Structural assembly | 1 Structural | False |
| CERS5 | 2 | hill_2022;li_2025 | M3 — Ciliary signaling | 4 Signaling Associated | False |
| CETN3 | 2 | hill_2022;li_2025 | M1 — Structural assembly | 6 Cytoskeleton ciliogenesis regulation | False |
| CFAP70 | 2 | hill_2022;li_2025 | M7 — Motile cilia (non-primary) | 7 Motile Cilia | False |
| CHMP4B | 2 | hill_2022;li_2025 | M4 — Turnover and maintenance | 5 Turnover/Maintenance | False |
| CKAP2L | 2 | hill_2022;li_2025 | M1 — Structural assembly | 6 Cytoskeleton ciliogenesis regulation | False |
| CKAP5 | 2 | hill_2022;li_2025 | M1 — Structural assembly | 6 Cytoskeleton ciliogenesis regulation | False |
| CLNS1A | 2 | hill_2022;li_2025 | M1 — Structural assembly | 6 Cytoskeleton ciliogenesis regulation | False |
| CLTA | 2 | hill_2022;li_2025 | M1 — Structural assembly | 6 Cytoskeleton ciliogenesis regulation | False |
| CLTC | 2 | hill_2022;li_2025 | M1 — Structural assembly | 1 Structural | False |
| CNTRL | 2 | hill_2022;li_2025 | M1 — Structural assembly | 1 Structural | False |
| COPS8 | 2 | hill_2022;li_2025 | M2 — Intraflagellar trafficking | 3 IFT/Trafficking/BBSome | False |
| CREB3 | 2 | hill_2022;li_2025 | M1 — Structural assembly | 1 Structural | False |
| CRK | 2 | hill_2022;li_2025 | M1 — Structural assembly | 1 Structural | False |
| CSNK1A1 | 2 | hill_2022;li_2025 | M3 — Ciliary signaling | 4 Signaling Associated | False |
| CSNK1D | 2 | hill_2022;li_2025 | M3 — Ciliary signaling | 4 Signaling Associated | False |
| CSNK2A1 | 2 | hill_2022;li_2025 | M3 — Ciliary signaling | 4 Signaling Associated | False |
| CTNND1 | 2 | hill_2022;li_2025 | M3 — Ciliary signaling | 4 Signaling Associated | False |
| CYFIP2 | 2 | hill_2022;li_2025 | M3 — Ciliary signaling | 4 Signaling Associated | False |
| DAAM1 | 2 | hill_2022;li_2025 | M1 — Structural assembly | 6 Cytoskeleton ciliogenesis regulation | False |
| DAPK3 | 2 | hill_2022;li_2025 | M2 — Intraflagellar trafficking | 3 IFT/Trafficking/BBSome | False |
| DCTN2 | 2 | hill_2022;li_2025 | M1 — Structural assembly | 6 Cytoskeleton ciliogenesis regulation | False |
| DCTN3 | 2 | hill_2022;li_2025 | M1 — Structural assembly | 6 Cytoskeleton ciliogenesis regulation | False |
| DDAH1 | 2 | de_soysa_2019;hill_2022 | M1 — Structural assembly | 1 Structural | False |
| DDX6 | 2 | hill_2022;li_2025 | M1 — Structural assembly | 1 Structural | False |
| DEGS1 | 2 | hill_2022;li_2025 | M3 — Ciliary signaling | 4 Signaling Associated | False |
| DHRS3 | 2 | hill_2022;li_2025 | M1 — Structural assembly | 1 Structural | False |
| DLD | 2 | hill_2022;li_2025 | M1 — Structural assembly | 1 Structural | False |
| DMD | 2 | hill_2022;soh_2021 | M1 — Structural assembly | 1 Structural | False |
| DNAJA1 | 2 | de_soysa_2019;hill_2022 | M1 — Structural assembly | 1 Structural | False |
| DNMBP | 2 | hill_2022;li_2025 | M3 — Ciliary signaling | 4 Signaling Associated | False |
| DPYSL2 | 2 | hill_2022;li_2025 | M1 — Structural assembly | 1 Structural | False |
| DYNC1H1 | 2 | hill_2022;li_2025 | M2 — Intraflagellar trafficking | 3 IFT/Trafficking/BBSome | False |
| DYNC1I2 | 2 | hill_2022;li_2025 | M7 — Motile cilia (non-primary) | 7 Motile Cilia | False |
| DYNLRB1 | 2 | hill_2022;li_2025 | M2 — Intraflagellar trafficking | 3 IFT/Trafficking/BBSome | False |
| E2F4 | 2 | hill_2022;li_2025 | M1 — Structural assembly | 6 Cytoskeleton ciliogenesis regulation | False |
| EFCAB1 | 2 | hill_2022;li_2025 | M3 — Ciliary signaling | 4 Signaling Associated | False |
| EFTUD2 | 2 | hill_2022;li_2025 | M1 — Structural assembly | 2 Assembly | False |
| EHD1 | 2 | hill_2022;li_2025 | M2 — Intraflagellar trafficking | 3 IFT/Trafficking/BBSome | False |
| EHD2 | 2 | hill_2022;li_2025 | M1 — Structural assembly | 1 Structural | False |
| EHD3 | 2 | hill_2022;lee_2023 | M2 — Intraflagellar trafficking | 3 IFT/Trafficking/BBSome | False |
| EIF5 | 2 | hill_2022;li_2025 | M2 — Intraflagellar trafficking | 3 IFT/Trafficking/BBSome | False |
| ENAH | 2 | hill_2022;li_2025 | M1 — Structural assembly | 1 Structural | False |
| EPB41L3 | 2 | hill_2022;li_2025 | M1 — Structural assembly | 1 Structural | False |
| EPS15 | 2 | hill_2022;li_2025 | M3 — Ciliary signaling | 4 Signaling Associated | False |
| EXOC6 | 2 | hill_2022;li_2025 | M2 — Intraflagellar trafficking | 3 IFT/Trafficking/BBSome | False |
| EXOC6B | 2 | hill_2022;li_2025 | M2 — Intraflagellar trafficking | 3 IFT/Trafficking/BBSome | False |
| FAM184A | 2 | hill_2022;li_2025 | M1 — Structural assembly | 1 Structural | False |
| FLAD1 | 2 | hill_2022;li_2025 | M2 — Intraflagellar trafficking | 3 IFT/Trafficking/BBSome | False |
| FLNA | 2 | hill_2022;li_2025 | M1 — Structural assembly | 6 Cytoskeleton ciliogenesis regulation | False |
| FNBP1L | 2 | hill_2022;li_2025 | M3 — Ciliary signaling | 4 Signaling Associated | False |
| FZD4 | 2 | hill_2022;li_2025 | M3 — Ciliary signaling | 4 Signaling Associated | False |
| GIGYF1 | 2 | hill_2022;li_2025 | M1 — Structural assembly | 6 Cytoskeleton ciliogenesis regulation | False |
| GLI3 | 2 | hill_2022;li_2025 | M3 — Ciliary signaling | 4 Signaling Associated | False |
| GLIS2 | 2 | hill_2022;li_2025 | M1 — Structural assembly | 1 Structural | False |
| GMNN | 2 | hill_2022;soh_2021 | M1 — Structural assembly | 1 Structural | False |
| GNAI2 | 2 | hill_2022;li_2025 | M1 — Structural assembly | 1 Structural | False |
| GNB1 | 2 | hill_2022;li_2025 | M3 — Ciliary signaling | 4 Signaling Associated | False |
| GNB5 | 2 | hill_2022;li_2025 | M3 — Ciliary signaling | 4 Signaling Associated | False |
| GOLGB1 | 2 | hill_2022;li_2025 | M2 — Intraflagellar trafficking | 3 IFT/Trafficking/BBSome | False |
| GPI | 2 | hill_2022;li_2025 | M1 — Structural assembly | 1 Structural | False |
| GPR183 | 2 | hill_2022;li_2025 | M3 — Ciliary signaling | 4 Signaling Associated | False |
| GPR22 | 2 | hill_2022;li_2025 | M3 — Ciliary signaling | 4 Signaling Associated | False |
| GRK4 | 2 | hill_2022;li_2025 | M3 — Ciliary signaling | 4 Signaling Associated | False |
| GSK3B | 2 | hill_2022;li_2025 | M3 — Ciliary signaling | 4 Signaling Associated | False |
| GSTA1 | 2 | lee_2023;soh_2021 | M1 — Structural assembly | 1 Structural | False |
| GSTM3 | 2 | hill_2022;li_2025 | M1 — Structural assembly | 1 Structural | False |
| GTSE1 | 2 | hill_2022;soh_2021 | M1 — Structural assembly | 1 Structural | False |
| HAVCR1 | 2 | hill_2022;li_2025 | M3 — Ciliary signaling | 4 Signaling Associated | False |
| HIF1A | 2 | hill_2022;li_2025 | M1 — Structural assembly | 2 Assembly | False |
| HRAS | 2 | hill_2022;li_2025 | M1 — Structural assembly | 1 Structural | False |
| HSP90B1 | 2 | hill_2022;li_2025 | M1 — Structural assembly | 1 Structural | False |
| HSPA8 | 2 | hill_2022;li_2025 | M1 — Structural assembly | 1 Structural | False |
| HSPB11 | 2 | hill_2022;li_2025 | M2 — Intraflagellar trafficking | 3 IFT/Trafficking/BBSome | False |
| HSPBP1 | 2 | hill_2022;li_2025 | M3 — Ciliary signaling | 4 Signaling Associated | False |
| HUWE1 | 2 | hill_2022;li_2025 | M4 — Turnover and maintenance | 5 Turnover/Maintenance | False |
| IFT140 | 2 | hill_2022;li_2025 | M7 — Motile cilia (non-primary) | 7 Motile Cilia | False |
| IFT57 | 2 | hill_2022;li_2025 | M7 — Motile cilia (non-primary) | 7 Motile Cilia | False |
| IFT74 | 2 | hill_2022;li_2025 | M7 — Motile cilia (non-primary) | 7 Motile Cilia | False |
| IGF1 | 2 | de_soysa_2019;hill_2022 | M3 — Ciliary signaling | 4 Signaling Associated | False |
| INSR | 2 | hill_2022;li_2025 | M1 — Structural assembly | 1 Structural | False |
| IPO5 | 2 | hill_2022;li_2025 | M1 — Structural assembly | 1 Structural | False |
| IRS1 | 2 | hill_2022;li_2025 | M3 — Ciliary signaling | 4 Signaling Associated | False |
| JADE1 | 2 | hill_2022;li_2025 | M1 — Structural assembly | 1 Structural | False |
| KCTD17 | 2 | hill_2022;li_2025 | M3 — Ciliary signaling | 4 Signaling Associated | False |
| KIAA1671 | 2 | hill_2022;li_2025 | M1 — Structural assembly | 6 Cytoskeleton ciliogenesis regulation | False |
| KIF11 | 2 | hill_2022;li_2025 | M1 — Structural assembly | 1 Structural | False |
| KIF13B | 2 | hill_2022;li_2025 | M2 — Intraflagellar trafficking | 3 IFT/Trafficking/BBSome | False |
| KIF14 | 2 | hill_2022;li_2025 | M2 — Intraflagellar trafficking | 3 IFT/Trafficking/BBSome | False |
| KIF5B | 2 | hill_2022;li_2025 | M2 — Intraflagellar trafficking | 3 IFT/Trafficking/BBSome | False |
| KIFC1 | 2 | hill_2022;li_2025 | M2 — Intraflagellar trafficking | 3 IFT/Trafficking/BBSome | False |
| KLC1 | 2 | hill_2022;li_2025 | M2 — Intraflagellar trafficking | 3 IFT/Trafficking/BBSome | False |
| KLHL4 | 2 | de_soysa_2019;hill_2022 | M1 — Structural assembly | 6 Cytoskeleton ciliogenesis regulation | False |
| KMT5B | 2 | hill_2022;li_2025 | M1 — Structural assembly | 1 Structural | False |
| LAMA5 | 2 | hill_2022;li_2025 | M1 — Structural assembly | 1 Structural | False |
| LDHB | 2 | hill_2022;li_2025 | M2 — Intraflagellar trafficking | 3 IFT/Trafficking/BBSome | False |
| LEPR | 2 | hill_2022;li_2025 | M3 — Ciliary signaling | 4 Signaling Associated | False |
| LGR4 | 2 | hill_2022;li_2025 | M3 — Ciliary signaling | 4 Signaling Associated | False |
| LIMA1 | 2 | hill_2022;li_2025 | M1 — Structural assembly | 6 Cytoskeleton ciliogenesis regulation | False |
| LRBA | 2 | hill_2022;li_2025 | M2 — Intraflagellar trafficking | 3 IFT/Trafficking/BBSome | False |
| LUZP1 | 2 | hill_2022;li_2025 | M1 — Structural assembly | 1 Structural | False |
| LZTFL1 | 2 | hill_2022;li_2025 | M2 — Intraflagellar trafficking | 3 IFT/Trafficking/BBSome | False |
| MAP1A | 2 | hill_2022;li_2025 | M1 — Structural assembly | 6 Cytoskeleton ciliogenesis regulation | False |
| MAP1LC3A | 2 | hill_2022;li_2025 | M1 — Structural assembly | 6 Cytoskeleton ciliogenesis regulation | False |
| MAP1LC3B | 2 | hill_2022;li_2025 | M1 — Structural assembly | 6 Cytoskeleton ciliogenesis regulation | False |
| MAP2K1 | 2 | hill_2022;li_2025 | M1 — Structural assembly | 1 Structural | False |
| MAP4 | 2 | hill_2022;zulibiya_2023 | M1 — Structural assembly | 6 Cytoskeleton ciliogenesis regulation | True |
| MAP7D3 | 2 | hill_2022;li_2025 | M1 — Structural assembly | 1 Structural | False |
| MAPK3 | 2 | hill_2022;li_2025 | M1 — Structural assembly | 6 Cytoskeleton ciliogenesis regulation | False |
| MAPK6 | 2 | hill_2022;li_2025 | M1 — Structural assembly | 6 Cytoskeleton ciliogenesis regulation | False |
| MAPKAPK2 | 2 | hill_2022;li_2025 | M1 — Structural assembly | 1 Structural | False |
| MAPRE1 | 2 | hill_2022;li_2025 | M1 — Structural assembly | 6 Cytoskeleton ciliogenesis regulation | False |
| MAPRE3 | 2 | hill_2022;li_2025 | M1 — Structural assembly | 6 Cytoskeleton ciliogenesis regulation | False |
| MARCKSL1 | 2 | hill_2022;lee_2023 | M1 — Structural assembly | 6 Cytoskeleton ciliogenesis regulation | False |
| MARK4 | 2 | hill_2022;li_2025 | M1 — Structural assembly | 6 Cytoskeleton ciliogenesis regulation | False |
| MCM2 | 2 | hill_2022;li_2025 | M4 — Turnover and maintenance | 5 Turnover/Maintenance | False |
| MCM7 | 2 | hill_2022;li_2025 | M1 — Structural assembly | 1 Structural | False |
| MERTK | 2 | hill_2022;li_2025 | M1 — Structural assembly | 1 Structural | False |
| MICAL1 | 2 | hill_2022;li_2025 | M1 — Structural assembly | 6 Cytoskeleton ciliogenesis regulation | False |
| MINDY3 | 2 | hill_2022;li_2025 | M1 — Structural assembly | 6 Cytoskeleton ciliogenesis regulation | False |
| MRPL11 | 2 | hill_2022;li_2025 | M1 — Structural assembly | 1 Structural | False |
| MRPS5 | 2 | hill_2022;li_2025 | M1 — Structural assembly | 1 Structural | False |
| MTOR | 2 | hill_2022;li_2025 | M3 — Ciliary signaling | 4 Signaling Associated | False |
| MYO7A | 2 | hill_2022;li_2025 | M1 — Structural assembly | 6 Cytoskeleton ciliogenesis regulation | False |
| MYOF | 2 | hill_2022;li_2025 | M1 — Structural assembly | 6 Cytoskeleton ciliogenesis regulation | False |
| MYRIP | 2 | hill_2022;zulibiya_2023 | M1 — Structural assembly | 6 Cytoskeleton ciliogenesis regulation | True |
| NAA15 | 2 | hill_2022;li_2025 | M1 — Structural assembly | 6 Cytoskeleton ciliogenesis regulation | False |
| NCAPD2 | 2 | hill_2022;li_2025 | M1 — Structural assembly | 1 Structural | False |
| NCBP2 | 2 | hill_2022;li_2025 | M1 — Structural assembly | 1 Structural | False |
| NEK2 | 2 | hill_2022;li_2025 | M4 — Turnover and maintenance | 5 Turnover/Maintenance | False |
| NFE2L2 | 2 | hill_2022;li_2025 | M3 — Ciliary signaling | 4 Signaling Associated | False |
| NGFR | 2 | hill_2022;lee_2023 | M3 — Ciliary signaling | 4 Signaling Associated | False |
| NINL | 2 | hill_2022;li_2025 | M1 — Structural assembly | 6 Cytoskeleton ciliogenesis regulation | False |
| NME3 | 2 | hill_2022;li_2025 | M1 — Structural assembly | 1 Structural | False |
| NUBP1 | 2 | hill_2022;li_2025 | M3 — Ciliary signaling | 4 Signaling Associated | False |
| NUDC | 2 | hill_2022;li_2025 | M1 — Structural assembly | 6 Cytoskeleton ciliogenesis regulation | False |
| NUP205 | 2 | hill_2022;li_2025 | M1 — Structural assembly | 1 Structural | False |
| NUP37 | 2 | hill_2022;li_2025 | M1 — Structural assembly | 1 Structural | False |
| NUP62 | 2 | hill_2022;li_2025 | M1 — Structural assembly | 1 Structural | False |
| NUP85 | 2 | hill_2022;li_2025 | M1 — Structural assembly | 1 Structural | False |
| NUP93 | 2 | hill_2022;li_2025 | M1 — Structural assembly | 1 Structural | False |
| OFD1 | 2 | hill_2022;li_2025 | M1 — Structural assembly | 6 Cytoskeleton ciliogenesis regulation | False |
| OSCP1 | 2 | hill_2022;li_2025 | M1 — Structural assembly | 1 Structural | False |
| PACSIN2 | 2 | hill_2022;li_2025 | M1 — Structural assembly | 1 Structural | False |
| PAFAH1B1 | 2 | hill_2022;li_2025 | M1 — Structural assembly | 6 Cytoskeleton ciliogenesis regulation | False |
| PARP4 | 2 | hill_2022;li_2025 | M1 — Structural assembly | 6 Cytoskeleton ciliogenesis regulation | False |
| PAWR | 2 | hill_2022;li_2025 | M1 — Structural assembly | 6 Cytoskeleton ciliogenesis regulation | False |
| PCNT | 2 | hill_2022;li_2025 | M1 — Structural assembly | 6 Cytoskeleton ciliogenesis regulation | False |
| PDE4B | 2 | hill_2022;li_2025 | M2 — Intraflagellar trafficking | 3 IFT/Trafficking/BBSome | False |
| PDE4D | 2 | hill_2022;zulibiya_2023 | M2 — Intraflagellar trafficking | 3 IFT/Trafficking/BBSome | True |
| PDE5A | 2 | hill_2022;li_2025 | M1 — Structural assembly | 1 Structural | False |
| PDIA6 | 2 | hill_2022;li_2025 | M1 — Structural assembly | 1 Structural | False |
| PELI1 | 2 | hill_2022;li_2025 | M1 — Structural assembly | 1 Structural | False |
| PFKM | 2 | hill_2022;li_2025 | M1 — Structural assembly | 1 Structural | False |
| PFN2 | 2 | hill_2022;li_2025 | M1 — Structural assembly | 1 Structural | False |
| PHTF1 | 2 | hill_2022;li_2025 | M1 — Structural assembly | 6 Cytoskeleton ciliogenesis regulation | False |
| PIK3C2A | 2 | hill_2022;li_2025 | M3 — Ciliary signaling | 4 Signaling Associated | False |
| PIN1 | 2 | hill_2022;li_2025 | M1 — Structural assembly | 6 Cytoskeleton ciliogenesis regulation | False |
| PKD2 | 2 | hill_2022;li_2025 | M3 — Ciliary signaling | 4 Signaling Associated | False |
| PKM | 2 | hill_2022;li_2025 | M1 — Structural assembly | 1 Structural | False |
| PLCB4 | 2 | hill_2022;zulibiya_2023 | M1 — Structural assembly | 6 Cytoskeleton ciliogenesis regulation | True |
| PLCG1 | 2 | hill_2022;li_2025 | M1 — Structural assembly | 1 Structural | False |
| POR | 2 | hill_2022;li_2025 | M1 — Structural assembly | 1 Structural | False |
| PPID | 2 | hill_2022;li_2025 | M2 — Intraflagellar trafficking | 3 IFT/Trafficking/BBSome | False |
| PPL | 2 | hill_2022;li_2025 | M1 — Structural assembly | 1 Structural | False |
| PPP2CB | 2 | hill_2022;li_2025 | M1 — Structural assembly | 1 Structural | False |
| PPP2R3B | 2 | hill_2022;li_2025 | M2 — Intraflagellar trafficking | 3 IFT/Trafficking/BBSome | False |
| PPP3R1 | 2 | hill_2022;li_2025 | M2 — Intraflagellar trafficking | 3 IFT/Trafficking/BBSome | False |
| PPP5C | 2 | hill_2022;li_2025 | M1 — Structural assembly | 1 Structural | False |
| PQBP1 | 2 | hill_2022;li_2025 | M3 — Ciliary signaling | 4 Signaling Associated | False |
| PRKAA1 | 2 | hill_2022;li_2025 | M3 — Ciliary signaling | 4 Signaling Associated | False |
| PRKAB1 | 2 | hill_2022;li_2025 | M1 — Structural assembly | 1 Structural | False |
| PRKACA | 2 | hill_2022;li_2025 | M3 — Ciliary signaling | 4 Signaling Associated | False |
| PRKAR1A | 2 | hill_2022;li_2025 | M3 — Ciliary signaling | 4 Signaling Associated | False |
| PRKAR2A | 2 | hill_2022;li_2025 | M3 — Ciliary signaling | 4 Signaling Associated | False |
| PRPF6 | 2 | hill_2022;li_2025 | M1 — Structural assembly | 1 Structural | False |
| PRPF8 | 2 | hill_2022;li_2025 | M1 — Structural assembly | 1 Structural | False |
| PSEN2 | 2 | hill_2022;li_2025 | M1 — Structural assembly | 1 Structural | False |
| PSMA6 | 2 | hill_2022;li_2025 | M1 — Structural assembly | 1 Structural | False |
| PSMB7 | 2 | hill_2022;li_2025 | M4 — Turnover and maintenance | 5 Turnover/Maintenance | False |
| PSMD5 | 2 | hill_2022;li_2025 | M1 — Structural assembly | 1 Structural | False |
| PSME3 | 2 | hill_2022;li_2025 | M1 — Structural assembly | 1 Structural | False |
| PTK2 | 2 | hill_2022;zulibiya_2023 | M1 — Structural assembly | 6 Cytoskeleton ciliogenesis regulation | True |
| PTPN13 | 2 | hill_2022;li_2025 | M1 — Structural assembly | 1 Structural | False |
| PTPRK | 2 | hill_2022;li_2025 | M3 — Ciliary signaling | 4 Signaling Associated | False |
| PTPRM | 2 | hill_2022;zulibiya_2023 | M3 — Ciliary signaling | 4 Signaling Associated | True |
| RAB10 | 2 | hill_2022;li_2025 | M2 — Intraflagellar trafficking | 3 IFT/Trafficking/BBSome | False |
| RAB11B | 2 | hill_2022;li_2025 | M1 — Structural assembly | 1 Structural | False |
| RAB34 | 2 | hill_2022;li_2025 | M2 — Intraflagellar trafficking | 3 IFT/Trafficking/BBSome | False |
| RAB6A | 2 | hill_2022;li_2025 | M1 — Structural assembly | 1 Structural | False |
| RAB7A | 2 | hill_2022;li_2025 | M4 — Turnover and maintenance | 5 Turnover/Maintenance | False |
| RAB8A | 2 | hill_2022;li_2025 | M2 — Intraflagellar trafficking | 3 IFT/Trafficking/BBSome | False |
| RABGAP1L | 2 | hill_2022;li_2025 | M3 — Ciliary signaling | 4 Signaling Associated | False |
| RAN | 2 | hill_2022;li_2025 | M2 — Intraflagellar trafficking | 3 IFT/Trafficking/BBSome | False |
| RANBP1 | 2 | hill_2022;soh_2021 | M3 — Ciliary signaling | 4 Signaling Associated | False |
| RAPGEF2 | 2 | hill_2022;li_2025 | M1 — Structural assembly | 1 Structural | False |
| RAPGEF4 | 2 | hill_2022;zulibiya_2023 | M1 — Structural assembly | 1 Structural | True |
| RB1 | 2 | hill_2022;li_2025 | M1 — Structural assembly | 1 Structural | False |
| RCAN2 | 2 | hill_2022;li_2025 | M1 — Structural assembly | 1 Structural | False |
| RHOU | 2 | hill_2022;li_2025 | M1 — Structural assembly | 6 Cytoskeleton ciliogenesis regulation | False |
| RP2 | 2 | hill_2022;li_2025 | M1 — Structural assembly | 1 Structural | False |
| RRM1 | 2 | hill_2022;li_2025 | M1 — Structural assembly | 1 Structural | False |
| RSL1D1 | 2 | hill_2022;li_2025 | M1 — Structural assembly | 1 Structural | False |
| RUVBL1 | 2 | hill_2022;li_2025 | M1 — Structural assembly | 2 Assembly | False |
| SEC63 | 2 | hill_2022;li_2025 | M1 — Structural assembly | 1 Structural | False |
| SF3A1 | 2 | hill_2022;li_2025 | M1 — Structural assembly | 1 Structural | False |
| SF3B2 | 2 | hill_2022;li_2025 | M1 — Structural assembly | 1 Structural | False |
| SH3D21 | 2 | hill_2022;li_2025 | M1 — Structural assembly | 1 Structural | False |
| SIAH2 | 2 | de_soysa_2019;hill_2022 | M4 — Turnover and maintenance | 5 Turnover/Maintenance | False |
| SIRT2 | 2 | hill_2022;li_2025 | M4 — Turnover and maintenance | 5 Turnover/Maintenance | False |
| SLC12A2 | 2 | hill_2022;li_2025 | M1 — Structural assembly | 1 Structural | False |
| SLC12A4 | 2 | hill_2022;li_2025 | M1 — Structural assembly | 1 Structural | False |
| SLC1A5 | 2 | hill_2022;li_2025 | M3 — Ciliary signaling | 4 Signaling Associated | False |
| SLC9A3R1 | 2 | de_soysa_2019;hill_2022 | M3 — Ciliary signaling | 4 Signaling Associated | False |
| SLC9B2 | 2 | hill_2022;li_2025 | M1 — Structural assembly | 1 Structural | False |
| SLIRP | 2 | hill_2022;li_2025 | M3 — Ciliary signaling | 4 Signaling Associated | False |
| SMAD3 | 2 | hill_2022;li_2025 | M1 — Structural assembly | 1 Structural | False |
| SMAD7 | 2 | hill_2022;li_2025 | M1 — Structural assembly | 1 Structural | False |
| SMG7 | 2 | hill_2022;li_2025 | M1 — Structural assembly | 1 Structural | False |
| SMO | 2 | hill_2022;li_2025 | M3 — Ciliary signaling | 4 Signaling Associated | False |
| SNAP23 | 2 | hill_2022;li_2025 | M3 — Ciliary signaling | 4 Signaling Associated | False |
| SNAP29 | 2 | hill_2022;li_2025 | M2 — Intraflagellar trafficking | 3 IFT/Trafficking/BBSome | False |
| SNRNP200 | 2 | hill_2022;li_2025 | M2 — Intraflagellar trafficking | 3 IFT/Trafficking/BBSome | False |
| SNTB2 | 2 | hill_2022;li_2025 | M1 — Structural assembly | 1 Structural | False |
| SNX10 | 2 | hill_2022;li_2025 | M2 — Intraflagellar trafficking | 3 IFT/Trafficking/BBSome | False |
| SORD | 2 | hill_2022;soh_2021 | M1 — Structural assembly | 1 Structural | False |
| SPAG1 | 2 | hill_2022;li_2025 | M7 — Motile cilia (non-primary) | 7 Motile Cilia | False |
| SPICE1 | 2 | hill_2022;li_2025 | M1 — Structural assembly | 1 Structural | False |
| SPTAN1 | 2 | hill_2022;li_2025 | M1 — Structural assembly | 6 Cytoskeleton ciliogenesis regulation | False |
| SQSTM1 | 2 | hill_2022;li_2025 | M4 — Turnover and maintenance | 5 Turnover/Maintenance | False |
| SRGAP2 | 2 | hill_2022;zulibiya_2023 | M1 — Structural assembly | 1 Structural | True |
| SSX2IP | 2 | hill_2022;li_2025 | M1 — Structural assembly | 6 Cytoskeleton ciliogenesis regulation | False |
| ST13 | 2 | hill_2022;li_2025 | M1 — Structural assembly | 1 Structural | False |
| STAG1 | 2 | hill_2022;li_2025 | M1 — Structural assembly | 6 Cytoskeleton ciliogenesis regulation | False |
| STX3 | 2 | hill_2022;li_2025 | M2 — Intraflagellar trafficking | 3 IFT/Trafficking/BBSome | False |
| SYNE1 | 2 | hill_2022;li_2025 | M1 — Structural assembly | 6 Cytoskeleton ciliogenesis regulation | False |
| TGFB3 | 2 | hill_2022;li_2025 | M1 — Structural assembly | 1 Structural | False |
| TOM1L2 | 2 | hill_2022;li_2025 | M1 — Structural assembly | 1 Structural | False |
| TP53BP1 | 2 | hill_2022;li_2025 | M3 — Ciliary signaling | 4 Signaling Associated | False |
| TRAPPC10 | 2 | hill_2022;li_2025 | M3 — Ciliary signaling | 4 Signaling Associated | False |
| TRAPPC3 | 2 | hill_2022;li_2025 | M3 — Ciliary signaling | 4 Signaling Associated | False |
| TRPV4 | 2 | hill_2022;li_2025 | M3 — Ciliary signaling | 4 Signaling Associated | False |
| TSC1 | 2 | hill_2022;li_2025 | M1 — Structural assembly | 1 Structural | False |
| TTK | 2 | hill_2022;li_2025 | M1 — Structural assembly | 2 Assembly | False |
| TUBA4A | 2 | hill_2022;li_2025 | M1 — Structural assembly | 1 Structural | False |
| TUBB1 | 2 | hill_2022;li_2025 | M1 — Structural assembly | 6 Cytoskeleton ciliogenesis regulation | False |
| TUBB2A | 2 | hill_2022;soh_2021 | M1 — Structural assembly | 1 Structural | False |
| TUBB2B | 2 | de_soysa_2019;soh_2021 | M1 — Structural assembly | 1 Structural | False |
| TUBGCP5 | 2 | hill_2022;li_2025 | M1 — Structural assembly | 6 Cytoskeleton ciliogenesis regulation | False |
| TXNDC15 | 2 | hill_2022;li_2025 | M1 — Structural assembly | 1 Structural | False |
| UBE2L3 | 2 | hill_2022;li_2025 | M1 — Structural assembly | 1 Structural | False |
| UGGT1 | 2 | hill_2022;li_2025 | M1 — Structural assembly | 1 Structural | False |
| USP14 | 2 | hill_2022;li_2025 | M4 — Turnover and maintenance | 5 Turnover/Maintenance | False |
| USP8 | 2 | hill_2022;li_2025 | M4 — Turnover and maintenance | 5 Turnover/Maintenance | False |
| USP9X | 2 | hill_2022;li_2025 | M4 — Turnover and maintenance | 5 Turnover/Maintenance | False |
| VCP | 2 | hill_2022;li_2025 | M3 — Ciliary signaling | 4 Signaling Associated | False |
| VDAC3 | 2 | hill_2022;li_2025 | M3 — Ciliary signaling | 4 Signaling Associated | False |
| VPS35 | 2 | hill_2022;li_2025 | M4 — Turnover and maintenance | 5 Turnover/Maintenance | False |
| VPS37B | 2 | hill_2022;li_2025 | M2 — Intraflagellar trafficking | 3 IFT/Trafficking/BBSome | False |
| VPS37C | 2 | hill_2022;li_2025 | M1 — Structural assembly | 1 Structural | False |
| WASF3 | 2 | hill_2022;li_2025 | M1 — Structural assembly | 6 Cytoskeleton ciliogenesis regulation | False |
| WNT5A | 2 | de_soysa_2019;hill_2022 | M3 — Ciliary signaling | 4 Signaling Associated | False |
| WWTR1 | 2 | hill_2022;soh_2021 | M1 — Structural assembly | 6 Cytoskeleton ciliogenesis regulation | False |
| XPO1 | 2 | hill_2022;li_2025 | M2 — Intraflagellar trafficking | 3 IFT/Trafficking/BBSome | False |
| YAP1 | 2 | hill_2022;li_2025 | M1 — Structural assembly | 6 Cytoskeleton ciliogenesis regulation | False |
| YWHAE | 2 | hill_2022;li_2025 | M1 — Structural assembly | 1 Structural | False |
| YWHAG | 2 | hill_2022;li_2025 | M1 — Structural assembly | 1 Structural | False |
| YWHAQ | 2 | hill_2022;li_2025 | M1 — Structural assembly | 1 Structural | False |
| ZFAND2B | 2 | hill_2022;li_2025 | M1 — Structural assembly | 1 Structural | False |
| ZNF330 | 2 | hill_2022;li_2025 | M1 — Structural assembly | 1 Structural | False |
| ZNF423 | 2 | hill_2022;li_2025 | M1 — Structural assembly | 6 Cytoskeleton ciliogenesis regulation | False |

**Table S3. 24-gene TOF endothelial anchor signature**

| **Gene** | **log2FC** | **FDR** | **Direction** | **Cell type** | **CCTP module** | **Cilia module** | **N datasets** | **Datasets** |
| --- | --- | --- | --- | --- | --- | --- | --- | --- |
| PTPRM | 2.60584905935914 | 0.0 | up | endothelial | M3 — Ciliary signaling | 4 Signaling Associated | 2 | hill_2022;zulibiya_2023 |
| ARL15 | 1.14306497870446 | 0.0 | up | endothelial | M1 — Structural assembly | 1 Structural | 2 | hill_2022;zulibiya_2023 |
| MAP4 | 1.04726748363596 | 0.0 | up | endothelial | M1 — Structural assembly | 6 Cytoskeleton ciliogenesis regulation | 2 | hill_2022;zulibiya_2023 |
| SMAD6 | 1.0207653310764 | 0.0 | up | endothelial | M1 — Structural assembly | 1 Structural | 3 | hill_2022;li_2025;zulibiya_2023 |
| AKT3 | 0.795593528459621 | 0.0 | up | endothelial | M1 — Structural assembly | 1 Structural | 2 | hill_2022;zulibiya_2023 |
| PTK2 | 0.730573094112915 | 0.0 | up | endothelial | M1 — Structural assembly | 6 Cytoskeleton ciliogenesis regulation | 2 | hill_2022;zulibiya_2023 |
| ABCC4 | 0.725726591690901 | 0.0 | up | endothelial | M3 — Ciliary signaling | 4 Signaling Associated | 3 | hill_2022;li_2025;zulibiya_2023 |
| ABLIM1 | 0.701385214044177 | 0.0 | up | endothelial | M1 — Structural assembly | 6 Cytoskeleton ciliogenesis regulation | 3 | hill_2022;li_2025;zulibiya_2023 |
| EGFR | 0.63822375397973 | 0.0 | up | endothelial | M3 — Ciliary signaling | 4 Signaling Associated | 3 | hill_2022;li_2025;zulibiya_2023 |
| SRGAP2 | 0.630609696955862 | 0.0 | up | endothelial | M1 — Structural assembly | 1 Structural | 2 | hill_2022;zulibiya_2023 |
| FGFR1 | 0.629842466514266 | 0.0 | up | endothelial | M3 — Ciliary signaling | 4 Signaling Associated | 3 | hill_2022;li_2025;zulibiya_2023 |
| EML1 | 0.539058954415753 | 0.0 | up | endothelial | M1 — Structural assembly | 6 Cytoskeleton ciliogenesis regulation | 3 | hill_2022;li_2025;zulibiya_2023 |
| NEDD9 | 0.519865481087982 | 0.0 | up | endothelial | M1 — Structural assembly | 1 Structural | 3 | hill_2022;li_2025;zulibiya_2023 |
| ABLIM3 | 0.515241821480401 | 8.60171121560572e-193 | up | endothelial | M1 — Structural assembly | 6 Cytoskeleton ciliogenesis regulation | 3 | hill_2022;li_2025;zulibiya_2023 |
| UTRN | 0.416823701670084 | 5.03838150924666e-109 | up | endothelial | M1 — Structural assembly | 1 Structural | 3 | hill_2022;li_2025;zulibiya_2023 |
| PLCB4 | 0.366464930241764 | 2.63518324064855e-98 | up | endothelial | M1 — Structural assembly | 6 Cytoskeleton ciliogenesis regulation | 2 | hill_2022;zulibiya_2023 |
| ACTN1 | 0.17088903386513 | 2.30594326839931e-16 |  | endothelial | M1 — Structural assembly | 6 Cytoskeleton ciliogenesis regulation | 4 | hill_2022;lee_2023;li_2025;zulibiya_2023 |
| PDE4D | 0.14006258810176 | 6.15233616436003e-96 |  | endothelial | M2 — Intraflagellar trafficking | 3 IFT/Trafficking/BBSome | 2 | hill_2022;zulibiya_2023 |
| TUBB6 | 0.0929218050359418 | 0.0001298466186209 |  | endothelial | M1 — Structural assembly | 6 Cytoskeleton ciliogenesis regulation | 4 | de_soysa_2019;hill_2022;li_2025;zulibiya_2023 |
| ACTN4 | 0.0766154402877569 | 3.91189448676534e-07 |  | endothelial | M1 — Structural assembly | 6 Cytoskeleton ciliogenesis regulation | 3 | hill_2022;li_2025;zulibiya_2023 |
| ADAMTS9 | -0.0126947105346682 | 8.707292114638469e-128 |  | endothelial | M1 — Structural assembly | 2 Assembly | 2 | hill_2022;zulibiya_2023 |
| MYRIP | -0.0345772450413923 | 9.17026719997544e-06 |  | endothelial | M1 — Structural assembly | 6 Cytoskeleton ciliogenesis regulation | 2 | hill_2022;zulibiya_2023 |
| NUAK1 | -0.200201949389101 | 1.99713292246483e-43 |  | endothelial | M1 — Structural assembly | 6 Cytoskeleton ciliogenesis regulation | 3 | hill_2022;li_2025;zulibiya_2023 |
| RAPGEF4 | -0.243274807532687 | 2.4104381905629397e-62 |  | endothelial | M1 — Structural assembly | 1 Structural | 2 | hill_2022;zulibiya_2023 |

**Table S4. Genome-wide Cross-Species Concordance Score (CSCS) — 396 genes**

| **gene** | **n_datasets** | **cctp_module** | **human_log2fc** | **human_direction** | **mouse_valid** | **concord_mouse** | **li_chamber_valid** | **concord_zebrafish** | **CSCS** | **conservation_class** |
| --- | --- | --- | --- | --- | --- | --- | --- | --- | --- | --- |
| AAMP | 2 | M1 — Structural assembly | 0.0292220840022796 |  | False |  | False |  |  | no_cross_species_data |
| ABCC3 | 2 | M1 — Structural assembly | 0.0045443566345526 |  | False |  | False |  |  | no_cross_species_data |
| ABCC4 | 3 | M3 — Ciliary signaling | 0.725726591690901 | up | False |  | False |  |  | no_cross_species_data |
| ABLIM1 | 3 | M1 — Structural assembly | 0.701385214044177 | up | False |  | True | 1.0 | 1.0 | partially_conserved |
| ABLIM3 | 3 | M1 — Structural assembly | 0.515241821480401 | up | False |  | False |  |  | no_cross_species_data |
| ACAA2 | 2 | M1 — Structural assembly | 0.0136134927491015 |  | False |  | False |  |  | no_cross_species_data |
| ACLY | 2 | M1 — Structural assembly | 0.0210254528742099 |  | False |  | False |  |  | no_cross_species_data |
| ACTN1 | 4 | M1 — Structural assembly | 0.17088903386513 |  | False |  | False | 1.0 | 1.0 | partially_conserved |
| ACTN2 | 4 | M1 — Structural assembly | 0.330075273164826 | up | True | 1.0 | False |  | 1.0 | partially_conserved |
| ACTN4 | 3 | M1 — Structural assembly | 0.0766154402877569 |  | False |  | True | 1.0 | 1.0 | partially_conserved |
| ACTR2 | 2 | M1 — Structural assembly | 0.181927241401045 |  | False |  | True | 1.0 | 1.0 | partially_conserved |
| ACTR3 | 2 | M1 — Structural assembly | 0.048887933517462 |  | False |  | False | 1.0 | 1.0 | partially_conserved |
| ACVR1B | 2 | M3 — Ciliary signaling | 0.0438590080970058 |  | False |  | True | 1.0 | 1.0 | partially_conserved |
| ADAM17 | 2 | M3 — Ciliary signaling | 0.19747136655895 |  | False |  | False |  |  | no_cross_species_data |
| ADAMTS9 | 2 | M1 — Structural assembly | -0.0126947105346682 |  | False |  | False |  |  | no_cross_species_data |
| ADCY6 | 2 | M3 — Ciliary signaling | 0.0156002441529056 |  | False |  | False | 1.0 | 1.0 | partially_conserved |
| AK1 | 4 | M1 — Structural assembly | -0.0157473006921815 |  | True | 0.0 | False |  | 0.0 | divergent |
| AK2 | 2 | M1 — Structural assembly | -0.0067021011812835 |  | False |  | True | 0.0 | 0.0 | divergent |
| AKT3 | 2 | M1 — Structural assembly | 0.795593528459621 | up | False |  | False |  |  | no_cross_species_data |
| ALDH1A1 | 2 | M1 — Structural assembly | 0.0023194428055555 |  | False |  | False |  |  | no_cross_species_data |
| ANO10 | 2 | M3 — Ciliary signaling | 0.0774460336626764 |  | False |  | False |  |  | no_cross_species_data |
| ANXA1 | 4 | M1 — Structural assembly | 0.0289539193748199 |  | False |  | True | 1.0 | 1.0 | partially_conserved |
| ANXA6 | 3 | M1 — Structural assembly | -0.0058125685910428 |  | True | 0.0 | False |  | 0.0 | divergent |
| APOO | 2 | M1 — Structural assembly | -0.0202450650243591 |  | False |  | False | 0.0 | 0.0 | divergent |
| APP | 3 | M1 — Structural assembly | 0.0645181259735599 |  | False |  | True | 1.0 | 1.0 | partially_conserved |
| ARHGAP1 | 2 | M1 — Structural assembly | -0.0082927583919849 |  | False |  | False | 0.0 | 0.0 | divergent |
| ARHGAP35 | 2 | M3 — Ciliary signaling | 0.173016348377652 |  | False |  | False | 1.0 | 1.0 | partially_conserved |
| ARL15 | 2 | M1 — Structural assembly | 1.14306497870446 | up | False |  | False |  |  | no_cross_species_data |
| ARL3 | 2 | M2 — Intraflagellar trafficking | 0.0301034106390366 |  | False |  | False |  |  | no_cross_species_data |
| ARPP19 | 2 | M1 — Structural assembly | 0.104787021399135 |  | False |  | False | 1.0 | 1.0 | partially_conserved |
| ASAP1 | 2 | M2 — Intraflagellar trafficking | 0.466562632514102 | up | False |  | True | 1.0 | 1.0 | partially_conserved |
| ATF3 | 2 | M1 — Structural assembly | 7.25117760209081e-05 |  | False |  | True | 1.0 | 1.0 | partially_conserved |
| ATF6 | 2 | M1 — Structural assembly | 0.141607677921537 |  | False |  | False |  |  | no_cross_species_data |
| ATG16L1 | 2 | M4 — Turnover and maintenance | 0.0166667941534418 |  | False |  | False |  |  | no_cross_species_data |
| ATG3 | 2 | M4 — Turnover and maintenance | 0.0098324462654174 |  | False |  | False |  |  | no_cross_species_data |
| ATG5 | 2 | M4 — Turnover and maintenance | 0.0307068169188451 |  | False |  | False |  |  | no_cross_species_data |
| ATP2A2 | 5 | M2 — Intraflagellar trafficking | 0.346550593544645 | up | True | 1.0 | True | 1.0 | 1.0 | fully_conserved |
| ATP2B4 | 2 | M2 — Intraflagellar trafficking | 0.0198195496597636 |  | False |  | True | 1.0 | 1.0 | partially_conserved |
| ATP6V0D1 | 2 | M2 — Intraflagellar trafficking | 0.0044678977040272 |  | False |  | False |  |  | no_cross_species_data |
| ATP6V1C1 | 2 | M2 — Intraflagellar trafficking | -0.0112942037002325 |  | False |  | False |  |  | no_cross_species_data |
| ATP6V1D | 2 | M2 — Intraflagellar trafficking | 0.0415003276504704 |  | False |  | False |  |  | no_cross_species_data |
| AURKA | 2 | M4 — Turnover and maintenance | -0.0005065950954586 |  | False |  | False | 0.0 | 0.0 | divergent |
| BAG2 | 2 | M1 — Structural assembly | 0.0005511756381257 |  | False |  | True | 1.0 | 1.0 | partially_conserved |
| BAG3 | 2 | M1 — Structural assembly | 0.0139408264336926 |  | False |  | False | 1.0 | 1.0 | partially_conserved |
| BCL3 | 2 | M1 — Structural assembly | 0.0655700685542861 |  | False |  | False |  |  | no_cross_species_data |
| BMP2 | 4 | M1 — Structural assembly | -0.000438391983927 |  | True | 1.0 | False |  | 1.0 | partially_conserved |
| BUB1B | 2 | M1 — Structural assembly | 0.0099948278561295 |  | False |  | False | 1.0 | 1.0 | partially_conserved |
| C7ORF57 | 2 | M1 — Structural assembly | -0.0016823703109708 |  | False |  | False |  |  | no_cross_species_data |
| CACNA1C | 2 | M3 — Ciliary signaling | 0.388549590206136 | up | False |  | False |  |  | no_cross_species_data |
| CAMK1D | 2 | M1 — Structural assembly | 0.0844015236738525 |  | False |  | False |  |  | no_cross_species_data |
| CAMK2A | 2 | M1 — Structural assembly | 0.0003925583873568 |  | False |  | False |  |  | no_cross_species_data |
| CAMK2B | 2 | M1 — Structural assembly | -0.0055899721111262 |  | False |  | False |  |  | no_cross_species_data |
| CAMK2G | 2 | M1 — Structural assembly | -0.056396643697895 |  | False |  | True | 0.0 | 0.0 | divergent |
| CAMK2N2 | 2 | M1 — Structural assembly | 0.000716075934644 |  | False |  | False |  |  | no_cross_species_data |
| CAV1 | 2 | M3 — Ciliary signaling | -0.372242392866626 | down | False |  | True | 0.0 | 0.0 | divergent |
| CCSAP | 2 | M1 — Structural assembly | 0.0050629913338469 |  | False |  | False | 1.0 | 1.0 | partially_conserved |
| CCT8 | 2 | M1 — Structural assembly | -0.0168930009480642 |  | False |  | True | 0.0 | 0.0 | divergent |
| CDC20 | 3 | M1 — Structural assembly | -0.0022827729102272 |  | False |  | False |  |  | no_cross_species_data |
| CDC42EP5 | 3 | M2 — Intraflagellar trafficking | 0.0004590128786271 |  | False |  | True | 1.0 | 1.0 | partially_conserved |
| CDH23 | 2 | M3 — Ciliary signaling | -0.02100871033431 |  | False |  | False |  |  | no_cross_species_data |
| CDK1 | 4 | M1 — Structural assembly | -0.000389247367336 |  | False |  | False | 0.0 | 0.0 | divergent |
| CDK2 | 2 | M1 — Structural assembly | 0.0050142913354392 |  | False |  | False |  |  | no_cross_species_data |
| CDK5RAP2 | 2 | M1 — Structural assembly | 0.0539246900501654 |  | False |  | False |  |  | no_cross_species_data |
| CDKN1B | 2 | M1 — Structural assembly | -0.0059335507657043 |  | False |  | True | 0.0 | 0.0 | divergent |
| CENPF | 4 | M1 — Structural assembly | 0.0033746113308182 |  | False |  | False | 1.0 | 1.0 | partially_conserved |
| CEP170 | 2 | M1 — Structural assembly | 0.115386675851285 |  | False |  | False |  |  | no_cross_species_data |
| CEP350 | 2 | M1 — Structural assembly | 0.223084614983515 |  | False |  | False |  |  | no_cross_species_data |
| CEP55 | 3 | M1 — Structural assembly | 0.0035219505324473 |  | False |  | False | 1.0 | 1.0 | partially_conserved |
| CEP70 | 2 | M1 — Structural assembly | 0.0081074024486792 |  | False |  | False |  |  | no_cross_species_data |
| CERS5 | 2 | M3 — Ciliary signaling | 0.0253303763398817 |  | False |  | False |  |  | no_cross_species_data |
| CETN3 | 2 | M1 — Structural assembly | 0.0001851327613886 |  | False |  | False | 1.0 | 1.0 | partially_conserved |
| CHMP4B | 2 | M4 — Turnover and maintenance | -0.0263329015332838 |  | False |  | True | 0.0 | 0.0 | divergent |
| CKAP2 | 3 | M1 — Structural assembly | -0.0030285979866617 |  | False |  | False |  |  | no_cross_species_data |
| CKAP2L | 2 | M1 — Structural assembly | 0.0068096927079901 |  | False |  | False | 1.0 | 1.0 | partially_conserved |
| CKAP5 | 2 | M1 — Structural assembly | 0.0102187102962619 |  | False |  | False | 1.0 | 1.0 | partially_conserved |
| CLNS1A | 2 | M1 — Structural assembly | 0.0670016483022871 |  | False |  | False | 1.0 | 1.0 | partially_conserved |
| CLTA | 2 | M1 — Structural assembly | 0.0184648595474598 |  | False |  | True | 1.0 | 1.0 | partially_conserved |
| CLTC | 2 | M1 — Structural assembly | 0.080400007559985 |  | False |  | True | 1.0 | 1.0 | partially_conserved |
| CNTRL | 2 | M1 — Structural assembly | 0.14004869766949 |  | False |  | False | 1.0 | 1.0 | partially_conserved |
| COPS8 | 2 | M2 — Intraflagellar trafficking | 0.0368000546151396 |  | False |  | False |  |  | no_cross_species_data |
| CREB3 | 2 | M1 — Structural assembly | 0.0111787212707612 |  | False |  | True | 1.0 | 1.0 | partially_conserved |
| CRK | 2 | M1 — Structural assembly | 0.237315499011135 |  | False |  | False | 1.0 | 1.0 | partially_conserved |
| CSNK1A1 | 2 | M3 — Ciliary signaling | 0.367633358270754 | up | False |  | True | 1.0 | 1.0 | partially_conserved |
| CSNK1D | 2 | M3 — Ciliary signaling | 0.125164649565534 |  | False |  | True | 1.0 | 1.0 | partially_conserved |
| CSNK2A1 | 2 | M3 — Ciliary signaling | 0.0342941851818202 |  | False |  | True | 1.0 | 1.0 | partially_conserved |
| CTNNB1 | 3 | M3 — Ciliary signaling | 0.0614204079697815 |  | False |  | True | 1.0 | 1.0 | partially_conserved |
| CTNND1 | 2 | M3 — Ciliary signaling | 0.0589018531570074 |  | False |  | False | 1.0 | 1.0 | partially_conserved |
| CYFIP2 | 2 | M3 — Ciliary signaling | -0.0015851168664466 |  | False |  | False | 0.0 | 0.0 | divergent |
| DAAM1 | 2 | M1 — Structural assembly | 0.207062540741538 |  | False |  | False | 1.0 | 1.0 | partially_conserved |
| DAPK3 | 2 | M2 — Intraflagellar trafficking | -0.0189210042875488 |  | False |  | False |  |  | no_cross_species_data |
| DCTN2 | 2 | M1 — Structural assembly | -0.0136811663182569 |  | False |  | False | 0.0 | 0.0 | divergent |
| DCTN3 | 2 | M1 — Structural assembly | -0.0007538348999374 |  | False |  | False |  |  | no_cross_species_data |
| DDAH1 | 2 | M1 — Structural assembly | 0.0051880251720509 |  | True | 0.0 | False |  | 0.0 | divergent |
| DDX6 | 2 | M1 — Structural assembly | 0.103925500478648 |  | False |  | False |  |  | no_cross_species_data |
| DEGS1 | 2 | M3 — Ciliary signaling | 0.0178784464695082 |  | False |  | True | 1.0 | 1.0 | partially_conserved |
| DHRS3 | 2 | M1 — Structural assembly | 0.226564720026852 |  | False |  | True | 1.0 | 1.0 | partially_conserved |
| DLD | 2 | M1 — Structural assembly | 0.0430458850069509 |  | False |  | False |  |  | no_cross_species_data |
| DMD | 2 | M1 — Structural assembly | 0.452606420030635 | up | False |  | False |  |  | no_cross_species_data |
| DNAJA1 | 2 | M1 — Structural assembly | 0.0093452809837797 |  | True | 1.0 | False |  | 1.0 | partially_conserved |
| DNMBP | 2 | M3 — Ciliary signaling | -0.0081280955204814 |  | False |  | False |  |  | no_cross_species_data |
| DPYSL2 | 2 | M1 — Structural assembly | 0.387434524089164 | up | False |  | False |  |  | no_cross_species_data |
| DYNC1H1 | 2 | M2 — Intraflagellar trafficking | 0.0910936872993415 |  | False |  | False |  |  | no_cross_species_data |
| DYNC1I2 | 2 | M7 — Motile cilia (non-primary) | -0.0069349416699698 |  | False |  | False | 0.0 | 0.0 | divergent |
| DYNLL1 | 3 | M2 — Intraflagellar trafficking | -0.0154475122880812 |  | False |  | True | 0.0 | 0.0 | divergent |
| DYNLRB1 | 2 | M2 — Intraflagellar trafficking | -0.0171286313975893 |  | False |  | True | 0.0 | 0.0 | divergent |
| E2F4 | 2 | M1 — Structural assembly | 0.0406623194899723 |  | False |  | False | 1.0 | 1.0 | partially_conserved |
| EEF1A1 | 3 | M1 — Structural assembly | -0.0241606451745477 |  | False |  | False |  |  | no_cross_species_data |
| EFTUD2 | 2 | M1 — Structural assembly | 0.0140145815769351 |  | False |  | False | 1.0 | 1.0 | partially_conserved |
| EGFR | 3 | M3 — Ciliary signaling | 0.63822375397973 | up | False |  | False |  |  | no_cross_species_data |
| EHD1 | 2 | M2 — Intraflagellar trafficking | -0.0089294813217911 |  | False |  | True | 0.0 | 0.0 | divergent |
| EHD2 | 2 | M1 — Structural assembly | -0.0102916129900438 |  | False |  | True | 0.0 | 0.0 | divergent |
| EHD3 | 2 | M2 — Intraflagellar trafficking | -0.0019051030156591 |  | False |  | False |  |  | no_cross_species_data |
| EHD4 | 3 | M1 — Structural assembly | 0.180104706919483 |  | True | 1.0 | True | 1.0 | 1.0 | fully_conserved |
| EIF5 | 2 | M2 — Intraflagellar trafficking | 0.0907455734850834 |  | False |  | False | 1.0 | 1.0 | partially_conserved |
| EML1 | 3 | M1 — Structural assembly | 0.539058954415753 | up | False |  | False |  |  | no_cross_species_data |
| ENAH | 2 | M1 — Structural assembly | -0.0562722952607013 |  | False |  | False |  |  | no_cross_species_data |
| EPB41L3 | 2 | M1 — Structural assembly | 0.0259532310477171 |  | False |  | False |  |  | no_cross_species_data |
| EPS15 | 2 | M3 — Ciliary signaling | 0.30149315448262 | up | False |  | False |  |  | no_cross_species_data |
| EXOC6 | 2 | M2 — Intraflagellar trafficking | 0.16658533948718 |  | False |  | False | 1.0 | 1.0 | partially_conserved |
| EXOC6B | 2 | M2 — Intraflagellar trafficking | 0.278559881262602 | up | False |  | False | 1.0 | 1.0 | partially_conserved |
| EZR | 4 | M1 — Structural assembly | 0.0479383966774952 |  | True | 1.0 | False |  | 1.0 | partially_conserved |
| FAM184A | 2 | M1 — Structural assembly | 0.0080489086803992 |  | False |  | False |  |  | no_cross_species_data |
| FBN2 | 4 | M2 — Intraflagellar trafficking | 0.0128371970670391 |  | True | 0.0 | False |  | 0.0 | divergent |
| FGFR1 | 3 | M3 — Ciliary signaling | 0.629842466514266 | up | False |  | False |  |  | no_cross_species_data |
| FKBP1A | 3 | M1 — Structural assembly | 0.180088461632269 |  | False |  | False |  |  | no_cross_species_data |
| FLAD1 | 2 | M2 — Intraflagellar trafficking | 0.01287636546598 |  | False |  | False |  |  | no_cross_species_data |
| FLNA | 2 | M1 — Structural assembly | -0.0036427381326986 |  | False |  | True | 0.0 | 0.0 | divergent |
| FNBP1L | 2 | M3 — Ciliary signaling | 0.149053573439975 |  | False |  | False | 1.0 | 1.0 | partially_conserved |
| FSTL1 | 4 | M1 — Structural assembly | 0.0256911934829629 |  | True | 0.0 | True | 1.0 | 0.5 | partially_conserved |
| FZD4 | 2 | M3 — Ciliary signaling | 0.236049685066267 |  | False |  | True | 1.0 | 1.0 | partially_conserved |
| GIGYF1 | 2 | M1 — Structural assembly | -0.0139296874653106 |  | False |  | False | 0.0 | 0.0 | divergent |
| GLI3 | 2 | M3 — Ciliary signaling | 0.0699812662178139 |  | False |  | False | 1.0 | 1.0 | partially_conserved |
| GLIS2 | 2 | M1 — Structural assembly | 0.0172387752664225 |  | False |  | False |  |  | no_cross_species_data |
| GMNN | 2 | M1 — Structural assembly | 0.0217521228051925 |  | False |  | False |  |  | no_cross_species_data |
| GNAI1 | 3 | M1 — Structural assembly | 0.0374039444300771 |  | True | 1.0 | True | 1.0 | 1.0 | fully_conserved |
| GNAI2 | 2 | M1 — Structural assembly | 0.121926264323302 |  | False |  | True | 1.0 | 1.0 | partially_conserved |
| GNB1 | 2 | M3 — Ciliary signaling | 0.249741740367248 |  | False |  | True | 1.0 | 1.0 | partially_conserved |
| GNB5 | 2 | M3 — Ciliary signaling | 0.004003613461358 |  | False |  | False |  |  | no_cross_species_data |
| GOLGB1 | 2 | M2 — Intraflagellar trafficking | 0.0844215784959105 |  | False |  | False |  |  | no_cross_species_data |
| GPI | 2 | M1 — Structural assembly | -0.0127813689952435 |  | False |  | False | 0.0 | 0.0 | divergent |
| GPR183 | 2 | M3 — Ciliary signaling | -0.0004830595979612 |  | False |  | False |  |  | no_cross_species_data |
| GPR22 | 2 | M3 — Ciliary signaling | 0.0027790544418206 |  | False |  | False |  |  | no_cross_species_data |
| GRK4 | 2 | M3 — Ciliary signaling | 0.0182402192864372 |  | False |  | True | 1.0 | 1.0 | partially_conserved |
| GSK3B | 2 | M3 — Ciliary signaling | 0.214666138179221 |  | False |  | False | 1.0 | 1.0 | partially_conserved |
| GSN | 3 | M1 — Structural assembly | -0.0828993381511629 |  | True | 0.0 | True | 0.0 | 0.0 | divergent |
| GSTM3 | 2 | M1 — Structural assembly | 0.0026641317272195 |  | True | 1.0 | False |  | 1.0 | partially_conserved |
| GTSE1 | 2 | M1 — Structural assembly | 0.0059059669570478 |  | False |  | False |  |  | no_cross_species_data |
| HAVCR1 | 2 | M3 — Ciliary signaling | -0.0007844726241696 |  | False |  | False |  |  | no_cross_species_data |
| HIF1A | 2 | M1 — Structural assembly | 0.0339550813010632 |  | False |  | True | 1.0 | 1.0 | partially_conserved |
| HK1 | 3 | M1 — Structural assembly | 0.199628226639594 |  | True | 1.0 | True | 1.0 | 1.0 | fully_conserved |
| HMGB2 | 4 | M1 — Structural assembly | -0.0132347208106177 |  | False |  | True | 0.0 | 0.0 | divergent |
| HRAS | 2 | M1 — Structural assembly | -0.0094093711143046 |  | False |  | False |  |  | no_cross_species_data |
| HSP90AA1 | 3 | M1 — Structural assembly | 0.0372143635897797 |  | False |  | True | 1.0 | 1.0 | partially_conserved |
| HSP90AB1 | 3 | M2 — Intraflagellar trafficking | 0.0020765772880883 |  | False |  | True | 1.0 | 1.0 | partially_conserved |
| HSP90B1 | 2 | M1 — Structural assembly | -0.0489372860825115 |  | False |  | True | 0.0 | 0.0 | divergent |
| HSPA8 | 2 | M1 — Structural assembly | 0.0112690296740197 |  | False |  | True | 1.0 | 1.0 | partially_conserved |
| HSPB11 | 2 | M2 — Intraflagellar trafficking | 0.0261877827813849 |  | False |  | False |  |  | no_cross_species_data |
| HSPBP1 | 2 | M3 — Ciliary signaling | 0.0058656695622689 |  | False |  | False |  |  | no_cross_species_data |
| HUWE1 | 2 | M4 — Turnover and maintenance | 0.073990393993644 |  | False |  | False | 1.0 | 1.0 | partially_conserved |
| IFT140 | 2 | M7 — Motile cilia (non-primary) | 0.363142872505541 | up | False |  | False | 1.0 | 1.0 | partially_conserved |
| IFT57 | 2 | M7 — Motile cilia (non-primary) | -0.0100879506472424 |  | False |  | False |  |  | no_cross_species_data |
| IFT74 | 2 | M7 — Motile cilia (non-primary) | 0.0311354687351575 |  | False |  | False | 1.0 | 1.0 | partially_conserved |
| IGF1 | 2 | M3 — Ciliary signaling | -0.0001609132345525 |  | True | 0.0 | False |  | 0.0 | divergent |
| INSR | 2 | M1 — Structural assembly | 0.476634883235535 | up | False |  | True | 1.0 | 1.0 | partially_conserved |
| IPO5 | 2 | M1 — Structural assembly | -0.0069374061536791 |  | False |  | True | 0.0 | 0.0 | divergent |
| IRS1 | 2 | M3 — Ciliary signaling | 0.001707332737882 |  | False |  | False |  |  | no_cross_species_data |
| JADE1 | 2 | M1 — Structural assembly | 0.0066192027290921 |  | False |  | False | 1.0 | 1.0 | partially_conserved |
| KCTD17 | 2 | M3 — Ciliary signaling | -0.0036896156234322 |  | False |  | False | 0.0 | 0.0 | divergent |
| KIAA1671 | 2 | M1 — Structural assembly | 0.30424244670705 | up | False |  | True | 1.0 | 1.0 | partially_conserved |
| KIF11 | 2 | M1 — Structural assembly | -0.0032318276277409 |  | False |  | False | 0.0 | 0.0 | divergent |
| KIF13B | 2 | M2 — Intraflagellar trafficking | 0.30692973326285 | up | False |  | True | 1.0 | 1.0 | partially_conserved |
| KIF14 | 2 | M2 — Intraflagellar trafficking | 0.0030299224091542 |  | False |  | False |  |  | no_cross_species_data |
| KIF20B | 3 | M1 — Structural assembly | 0.0095974732312059 |  | False |  | False | 1.0 | 1.0 | partially_conserved |
| KIF5B | 2 | M2 — Intraflagellar trafficking | 0.0955799246147118 |  | False |  | False | 1.0 | 1.0 | partially_conserved |
| KIFC1 | 2 | M2 — Intraflagellar trafficking | 0.0004383871385099 |  | False |  | False | 1.0 | 1.0 | partially_conserved |
| KLC1 | 2 | M2 — Intraflagellar trafficking | 0.0380243081513358 |  | False |  | False |  |  | no_cross_species_data |
| KLHL4 | 2 | M1 — Structural assembly | -0.0007892289119822 |  | True | 0.0 | False |  | 0.0 | divergent |
| LAMA5 | 2 | M1 — Structural assembly | -0.0692364122021821 |  | False |  | False |  |  | no_cross_species_data |
| LDHB | 2 | M2 — Intraflagellar trafficking | 0.0410396737518625 |  | False |  | False |  |  | no_cross_species_data |
| LEPR | 2 | M3 — Ciliary signaling | 0.0258327971486221 |  | False |  | False |  |  | no_cross_species_data |
| LGR4 | 2 | M3 — Ciliary signaling | 0.0086994541866499 |  | False |  | False |  |  | no_cross_species_data |
| LIMA1 | 2 | M1 — Structural assembly | -0.13889411971466 |  | False |  | True | 0.0 | 0.0 | divergent |
| LRBA | 2 | M2 — Intraflagellar trafficking | 0.241180194303081 |  | False |  | True | 1.0 | 1.0 | partially_conserved |
| LRRC49 | 3 | M1 — Structural assembly | 0.0171709760184 |  | False |  | False |  |  | no_cross_species_data |
| LUZP1 | 2 | M1 — Structural assembly | 0.110996274301718 |  | False |  | True | 1.0 | 1.0 | partially_conserved |
| LZTFL1 | 2 | M2 — Intraflagellar trafficking | 0.008384311598713 |  | False |  | False | 1.0 | 1.0 | partially_conserved |
| MAP1A | 2 | M1 — Structural assembly | -0.0042648903329636 |  | False |  | False |  |  | no_cross_species_data |
| MAP1LC3A | 2 | M1 — Structural assembly | 0.0154703227303719 |  | False |  | False |  |  | no_cross_species_data |
| MAP1LC3B | 2 | M1 — Structural assembly | 0.0017606407263929 |  | False |  | False | 1.0 | 1.0 | partially_conserved |
| MAP2K1 | 2 | M1 — Structural assembly | 0.0750053543646858 |  | False |  | False | 1.0 | 1.0 | partially_conserved |
| MAP4 | 2 | M1 — Structural assembly | 1.04726748363596 | up | False |  | False |  |  | no_cross_species_data |
| MAP7D3 | 2 | M1 — Structural assembly | -0.0059176475842726 |  | False |  | False | 0.0 | 0.0 | divergent |
| MAPK3 | 2 | M1 — Structural assembly | 0.0067449674599589 |  | False |  | False | 1.0 | 1.0 | partially_conserved |
| MAPK6 | 2 | M1 — Structural assembly | 0.0516899016775536 |  | False |  | False |  |  | no_cross_species_data |
| MAPKAPK2 | 2 | M1 — Structural assembly | 0.108859240837478 |  | False |  | False |  |  | no_cross_species_data |
| MAPRE1 | 2 | M1 — Structural assembly | 0.0446440369191037 |  | False |  | True | 1.0 | 1.0 | partially_conserved |
| MAPRE3 | 2 | M1 — Structural assembly | -0.0089599443187815 |  | False |  | False |  |  | no_cross_species_data |
| MARCKS | 4 | M1 — Structural assembly | -0.0476311523532823 |  | True | 0.0 | False |  | 0.0 | divergent |
| MARCKSL1 | 2 | M1 — Structural assembly | 0.0115704235220569 |  | False |  | False |  |  | no_cross_species_data |
| MARK4 | 2 | M1 — Structural assembly | 0.10855261333064 |  | False |  | False |  |  | no_cross_species_data |
| MCM2 | 2 | M4 — Turnover and maintenance | -0.0005301637080999 |  | False |  | False | 0.0 | 0.0 | divergent |
| MCM7 | 2 | M1 — Structural assembly | 2.64823387592927e-05 |  | False |  | False | 1.0 | 1.0 | partially_conserved |
| MERTK | 2 | M1 — Structural assembly | 0.0436522065719261 |  | False |  | False |  |  | no_cross_species_data |
| MICAL1 | 2 | M1 — Structural assembly | -0.0096254196323765 |  | False |  | False |  |  | no_cross_species_data |
| MRPL11 | 2 | M1 — Structural assembly | -0.0049098548101031 |  | False |  | False |  |  | no_cross_species_data |
| MRPS5 | 2 | M1 — Structural assembly | 0.0021023786518039 |  | False |  | False |  |  | no_cross_species_data |
| MTOR | 2 | M3 — Ciliary signaling | 0.0512609334467966 |  | False |  | False |  |  | no_cross_species_data |
| MYH10 | 5 | M1 — Structural assembly | 0.142471351482861 |  | True | 1.0 | True | 1.0 | 1.0 | fully_conserved |
| MYO7A | 2 | M1 — Structural assembly | -0.0023979338477646 |  | False |  | False | 0.0 | 0.0 | divergent |
| MYOF | 2 | M1 — Structural assembly | 0.0293338540848702 |  | False |  | True | 1.0 | 1.0 | partially_conserved |
| MYRIP | 2 | M1 — Structural assembly | -0.0345772450413923 |  | False |  | False |  |  | no_cross_species_data |
| NAA15 | 2 | M1 — Structural assembly | 0.119319633113061 |  | False |  | False |  |  | no_cross_species_data |
| NCAPD2 | 2 | M1 — Structural assembly | -0.0095618457533948 |  | False |  | False | 0.0 | 0.0 | divergent |
| NCBP2 | 2 | M1 — Structural assembly | -0.0062012879817235 |  | False |  | False |  |  | no_cross_species_data |
| NEDD9 | 3 | M1 — Structural assembly | 0.519865481087982 | up | False |  | False |  |  | no_cross_species_data |
| NFE2L2 | 2 | M3 — Ciliary signaling | 0.140977655840188 |  | False |  | True | 1.0 | 1.0 | partially_conserved |
| NINL | 2 | M1 — Structural assembly | 0.0598472999654549 |  | False |  | False |  |  | no_cross_species_data |
| NME3 | 2 | M1 — Structural assembly | -0.0633804612862274 |  | False |  | False | 0.0 | 0.0 | divergent |
| NOTCH1 | 3 | M3 — Ciliary signaling | 0.0408065383293413 |  | True | 1.0 | True | 1.0 | 1.0 | fully_conserved |
| NUAK1 | 3 | M1 — Structural assembly | -0.200201949389101 |  | False |  | False |  |  | no_cross_species_data |
| NUBP1 | 2 | M3 — Ciliary signaling | 0.0001417550142284 |  | False |  | False |  |  | no_cross_species_data |
| NUDC | 2 | M1 — Structural assembly | -0.0201259193660561 |  | False |  | False |  |  | no_cross_species_data |
| NUP205 | 2 | M1 — Structural assembly | -0.0151464343017527 |  | False |  | False |  |  | no_cross_species_data |
| NUP37 | 2 | M1 — Structural assembly | -0.0135381289347267 |  | False |  | False | 0.0 | 0.0 | divergent |
| NUP85 | 2 | M1 — Structural assembly | 0.0111649128351876 |  | False |  | False |  |  | no_cross_species_data |
| NUP93 | 2 | M1 — Structural assembly | 0.0241329246703787 |  | False |  | False | 1.0 | 1.0 | partially_conserved |
| OFD1 | 2 | M1 — Structural assembly | 0.028221094102155 |  | False |  | False | 1.0 | 1.0 | partially_conserved |
| OSCP1 | 2 | M1 — Structural assembly | 0.003829034074744 |  | False |  | False |  |  | no_cross_species_data |
| PACSIN2 | 2 | M1 — Structural assembly | 0.408883620939517 | up | False |  | True | 1.0 | 1.0 | partially_conserved |
| PAFAH1B1 | 2 | M1 — Structural assembly | 0.292357436066192 | up | False |  | True | 1.0 | 1.0 | partially_conserved |
| PARP4 | 2 | M1 — Structural assembly | 0.029795957850242 |  | False |  | False |  |  | no_cross_species_data |
| PAWR | 2 | M1 — Structural assembly | -0.0216344642732712 |  | False |  | False | 0.0 | 0.0 | divergent |
| PCNT | 2 | M1 — Structural assembly | 0.0084573349507214 |  | False |  | False | 1.0 | 1.0 | partially_conserved |
| PDE4B | 2 | M2 — Intraflagellar trafficking | 0.0161994186253957 |  | False |  | True | 1.0 | 1.0 | partially_conserved |
| PDE4D | 2 | M2 — Intraflagellar trafficking | 0.14006258810176 |  | False |  | False |  |  | no_cross_species_data |
| PDE5A | 2 | M1 — Structural assembly | -0.0367054447841953 |  | False |  | False |  |  | no_cross_species_data |
| PDGFRA | 3 | M3 — Ciliary signaling | 0.0031133158994183 |  | False |  | True | 1.0 | 1.0 | partially_conserved |
| PDIA6 | 2 | M1 — Structural assembly | -0.0293929899830167 |  | False |  | False | 0.0 | 0.0 | divergent |
| PELI1 | 2 | M1 — Structural assembly | 0.16500201394494 |  | False |  | False |  |  | no_cross_species_data |
| PFKM | 2 | M1 — Structural assembly | 0.0054667800309426 |  | False |  | False |  |  | no_cross_species_data |
| PFN2 | 2 | M1 — Structural assembly | -0.0006769999273263 |  | False |  | True | 0.0 | 0.0 | divergent |
| PHGDH | 3 | M2 — Intraflagellar trafficking | -0.0008998062267426 |  | True | 0.0 | False |  | 0.0 | divergent |
| PHTF1 | 2 | M1 — Structural assembly | 0.0191107996244089 |  | False |  | False |  |  | no_cross_species_data |
| PIK3C2A | 2 | M3 — Ciliary signaling | 0.372179401067464 | up | False |  | True | 1.0 | 1.0 | partially_conserved |
| PIN1 | 2 | M1 — Structural assembly | -0.0108618610870373 |  | False |  | False | 0.0 | 0.0 | divergent |
| PKD2 | 2 | M3 — Ciliary signaling | 0.201159641189724 |  | False |  | False |  |  | no_cross_species_data |
| PKIG | 4 | M3 — Ciliary signaling | 0.0831691752688076 |  | False |  | True | 1.0 | 1.0 | partially_conserved |
| PKM | 2 | M1 — Structural assembly | 0.0372431107581942 |  | False |  | True | 1.0 | 1.0 | partially_conserved |
| PLCB4 | 2 | M1 — Structural assembly | 0.366464930241764 | up | False |  | False |  |  | no_cross_species_data |
| PLCG1 | 2 | M1 — Structural assembly | 0.073331973505022 |  | False |  | True | 1.0 | 1.0 | partially_conserved |
| PLK1 | 3 | M4 — Turnover and maintenance | -0.0031528602305767 |  | False |  | False | 0.0 | 0.0 | divergent |
| POR | 2 | M1 — Structural assembly | -0.0133696008486967 |  | False |  | False |  |  | no_cross_species_data |
| PPID | 2 | M2 — Intraflagellar trafficking | -0.0129939384250211 |  | False |  | False |  |  | no_cross_species_data |
| PPL | 2 | M1 — Structural assembly | -0.0430451498801835 |  | False |  | False |  |  | no_cross_species_data |
| PPP2CB | 2 | M1 — Structural assembly | 0.170867562888522 |  | False |  | True | 1.0 | 1.0 | partially_conserved |
| PPP2R3B | 2 | M2 — Intraflagellar trafficking | -0.0071701384966557 |  | False |  | False |  |  | no_cross_species_data |
| PPP5C | 2 | M1 — Structural assembly | 0.107587305447411 |  | False |  | False |  |  | no_cross_species_data |
| PQBP1 | 2 | M3 — Ciliary signaling | -0.0046719091593368 |  | False |  | False |  |  | no_cross_species_data |
| PRDX1 | 3 | M1 — Structural assembly | -1.9197826936667e-05 |  | False |  | False | 0.0 | 0.0 | divergent |
| PRKAA1 | 2 | M3 — Ciliary signaling | 0.067764583830162 |  | False |  | False |  |  | no_cross_species_data |
| PRKAB1 | 2 | M1 — Structural assembly | 0.0285122297185694 |  | False |  | False | 1.0 | 1.0 | partially_conserved |
| PRKACA | 2 | M3 — Ciliary signaling | 0.0462316864604255 |  | False |  | False | 1.0 | 1.0 | partially_conserved |
| PRKAR1A | 2 | M3 — Ciliary signaling | 0.0680372333598559 |  | False |  | True | 1.0 | 1.0 | partially_conserved |
| PRKAR2A | 2 | M3 — Ciliary signaling | 0.311694977458052 | up | False |  | True | 1.0 | 1.0 | partially_conserved |
| PRPF6 | 2 | M1 — Structural assembly | -0.0150683157420383 |  | False |  | False |  |  | no_cross_species_data |
| PRPF8 | 2 | M1 — Structural assembly | 0.0303239702809987 |  | False |  | True | 1.0 | 1.0 | partially_conserved |
| PSEN2 | 2 | M1 — Structural assembly | 0.0003678782142052 |  | False |  | False |  |  | no_cross_species_data |
| PSMA6 | 2 | M1 — Structural assembly | 0.0014316394714378 |  | False |  | False | 1.0 | 1.0 | partially_conserved |
| PSMB7 | 2 | M4 — Turnover and maintenance | 0.0360988779249622 |  | False |  | True | 1.0 | 1.0 | partially_conserved |
| PSMC2 | 3 | M4 — Turnover and maintenance | 0.0104947200324342 |  | True | 1.0 | False | 1.0 | 1.0 | fully_conserved |
| PSMC4 | 3 | M1 — Structural assembly | 0.0041222828875653 |  | True | 1.0 | False | 1.0 | 1.0 | fully_conserved |
| PSMC5 | 3 | M4 — Turnover and maintenance | -0.0052338089435366 |  | True | 0.0 | False | 0.0 | 0.0 | divergent |
| PSMD5 | 2 | M1 — Structural assembly | 0.0356100478561311 |  | False |  | False |  |  | no_cross_species_data |
| PSME3 | 2 | M1 — Structural assembly | -0.0083514983420593 |  | False |  | False |  |  | no_cross_species_data |
| PTK2 | 2 | M1 — Structural assembly | 0.730573094112915 | up | False |  | False |  |  | no_cross_species_data |
| PTPN13 | 2 | M1 — Structural assembly | 0.0002998533834885 |  | False |  | False |  |  | no_cross_species_data |
| PTPRK | 2 | M3 — Ciliary signaling | 0.205407997578776 |  | False |  | False |  |  | no_cross_species_data |
| PTPRM | 2 | M3 — Ciliary signaling | 2.60584905935914 | up | False |  | False |  |  | no_cross_species_data |
| RAB10 | 2 | M2 — Intraflagellar trafficking | 0.27408252482133 | up | False |  | True | 1.0 | 1.0 | partially_conserved |
| RAB11A | 4 | M4 — Turnover and maintenance | 0.0693083026135304 |  | True | 1.0 | True | 1.0 | 1.0 | fully_conserved |
| RAB11B | 2 | M1 — Structural assembly | 0.0087272216702876 |  | False |  | False | 1.0 | 1.0 | partially_conserved |
| RAB34 | 2 | M2 — Intraflagellar trafficking | 0.0066804603480276 |  | False |  | False |  |  | no_cross_species_data |
| RAB6A | 2 | M1 — Structural assembly | 0.288863405813625 | up | False |  | False |  |  | no_cross_species_data |
| RAB7A | 2 | M4 — Turnover and maintenance | 0.209808826286425 |  | False |  | False |  |  | no_cross_species_data |
| RAB8A | 2 | M2 — Intraflagellar trafficking | -0.0039640939045352 |  | False |  | False | 0.0 | 0.0 | divergent |
| RABGAP1L | 2 | M3 — Ciliary signaling | 0.490895693120184 | up | False |  | True | 1.0 | 1.0 | partially_conserved |
| RAN | 2 | M2 — Intraflagellar trafficking | -0.0015238899986034 |  | False |  | True | 0.0 | 0.0 | divergent |
| RANBP1 | 2 | M3 — Ciliary signaling | 0.0056364372846914 |  | False |  | False |  |  | no_cross_species_data |
| RAPGEF2 | 2 | M1 — Structural assembly | 0.266891241805151 | up | False |  | True | 1.0 | 1.0 | partially_conserved |
| RAPGEF4 | 2 | M1 — Structural assembly | -0.243274807532687 |  | False |  | False |  |  | no_cross_species_data |
| RB1 | 2 | M1 — Structural assembly | 0.140541129271059 |  | False |  | False |  |  | no_cross_species_data |
| RCAN2 | 2 | M1 — Structural assembly | 0.301894844121678 | up | False |  | False |  |  | no_cross_species_data |
| RCSD1 | 3 | M1 — Structural assembly | 0.0355607773302905 |  | True | 1.0 | False | 1.0 | 1.0 | fully_conserved |
| RHOU | 2 | M1 — Structural assembly | 0.0155438050636965 |  | False |  | False | 1.0 | 1.0 | partially_conserved |
| RP2 | 2 | M1 — Structural assembly | 0.0230784897865866 |  | False |  | False |  |  | no_cross_species_data |
| RRAD | 3 | M1 — Structural assembly | 0.0002766798760938 |  | True | 1.0 | False |  | 1.0 | partially_conserved |
| RRM1 | 2 | M1 — Structural assembly | 0.0042539550949495 |  | False |  | False |  |  | no_cross_species_data |
| RSL1D1 | 2 | M1 — Structural assembly | 0.017714740685224 |  | False |  | True | 1.0 | 1.0 | partially_conserved |
| RUVBL1 | 2 | M1 — Structural assembly | -0.0466227754134749 |  | False |  | False |  |  | no_cross_species_data |
| SCCPDH | 3 | M1 — Structural assembly | 0.013932176036431 |  | True | 1.0 | False |  | 1.0 | partially_conserved |
| SEC63 | 2 | M1 — Structural assembly | 0.0639826393723355 |  | False |  | True | 1.0 | 1.0 | partially_conserved |
| SF3A1 | 2 | M1 — Structural assembly | 0.0243054974214706 |  | False |  | True | 1.0 | 1.0 | partially_conserved |
| SF3B2 | 2 | M1 — Structural assembly | -0.0044018189339804 |  | False |  | True | 0.0 | 0.0 | divergent |
| SH3D21 | 2 | M1 — Structural assembly | -0.0072808330686072 |  | False |  | False | 0.0 | 0.0 | divergent |
| SIAH2 | 2 | M4 — Turnover and maintenance | -0.0122955424042448 |  | True | 0.0 | False |  | 0.0 | divergent |
| SIRT2 | 2 | M4 — Turnover and maintenance | 0.0037343025104854 |  | False |  | False |  |  | no_cross_species_data |
| SKP1 | 3 | M1 — Structural assembly | 0.0447395865420842 |  | False |  | False |  |  | no_cross_species_data |
| SLC12A2 | 2 | M1 — Structural assembly | -0.0691261241230044 |  | False |  | True | 0.0 | 0.0 | divergent |
| SLC12A4 | 2 | M1 — Structural assembly | -0.022005007349087 |  | False |  | True | 0.0 | 0.0 | divergent |
| SLC1A5 | 2 | M3 — Ciliary signaling | -0.0010540287414095 |  | False |  | False | 0.0 | 0.0 | divergent |
| SLC9A3R1 | 2 | M3 — Ciliary signaling | 0.0098773661569132 |  | True | 1.0 | False |  | 1.0 | partially_conserved |
| SLC9B2 | 2 | M1 — Structural assembly | -0.0446236926870786 |  | False |  | False |  |  | no_cross_species_data |
| SLIRP | 2 | M3 — Ciliary signaling | -0.0337984221427849 |  | False |  | False | 0.0 | 0.0 | divergent |
| SMAD3 | 2 | M1 — Structural assembly | 0.182027557588808 |  | False |  | True | 1.0 | 1.0 | partially_conserved |
| SMAD6 | 3 | M1 — Structural assembly | 1.0207653310764 | up | False |  | False |  |  | no_cross_species_data |
| SMAD7 | 2 | M1 — Structural assembly | 0.136759956079846 |  | False |  | True | 1.0 | 1.0 | partially_conserved |
| SMG7 | 2 | M1 — Structural assembly | 0.0942649114207898 |  | False |  | False |  |  | no_cross_species_data |
| SMO | 2 | M3 — Ciliary signaling | 0.0009866550567864 |  | False |  | False |  |  | no_cross_species_data |
| SNAP23 | 2 | M3 — Ciliary signaling | 0.20896653877264 |  | False |  | False |  |  | no_cross_species_data |
| SNAP29 | 2 | M2 — Intraflagellar trafficking | 0.0177136283400276 |  | False |  | False | 1.0 | 1.0 | partially_conserved |
| SNRNP200 | 2 | M2 — Intraflagellar trafficking | 0.0111523690736453 |  | False |  | False | 1.0 | 1.0 | partially_conserved |
| SNTB2 | 2 | M1 — Structural assembly | 0.0413185428451307 |  | False |  | True | 1.0 | 1.0 | partially_conserved |
| SNX10 | 2 | M2 — Intraflagellar trafficking | -0.0055022766684742 |  | False |  | False |  |  | no_cross_species_data |
| SORD | 2 | M1 — Structural assembly | -0.0020443591248048 |  | False |  | False |  |  | no_cross_species_data |
| SPAG1 | 2 | M7 — Motile cilia (non-primary) | 0.0186290394166654 |  | False |  | False |  |  | no_cross_species_data |
| SPICE1 | 2 | M1 — Structural assembly | 0.0132472912479848 |  | False |  | False |  |  | no_cross_species_data |
| SPTAN1 | 2 | M1 — Structural assembly | 0.0912783500092595 |  | False |  | True | 1.0 | 1.0 | partially_conserved |
| SQSTM1 | 2 | M4 — Turnover and maintenance | 0.0903411792684339 |  | False |  | True | 1.0 | 1.0 | partially_conserved |
| SRGAP2 | 2 | M1 — Structural assembly | 0.630609696955862 | up | False |  | False |  |  | no_cross_species_data |
| SSX2IP | 2 | M1 — Structural assembly | 0.0002998533834885 |  | False |  | False |  |  | no_cross_species_data |
| ST13 | 2 | M1 — Structural assembly | -0.021993035432778 |  | False |  | False |  |  | no_cross_species_data |
| STAG1 | 2 | M1 — Structural assembly | 0.350068554638371 | up | False |  | False | 1.0 | 1.0 | partially_conserved |
| STX3 | 2 | M2 — Intraflagellar trafficking | 0.110430670973367 |  | False |  | True | 1.0 | 1.0 | partially_conserved |
| SYNE1 | 2 | M1 — Structural assembly | -0.220409694338377 |  | False |  | False |  |  | no_cross_species_data |
| TACC3 | 3 | M1 — Structural assembly | -0.0034560786864245 |  | False |  | False | 0.0 | 0.0 | divergent |
| TAGLN2 | 4 | M1 — Structural assembly | -0.0521119060559703 |  | True | 0.0 | False |  | 0.0 | divergent |
| TEK | 3 | M3 — Ciliary signaling | 0.0981219903593969 |  | True | 1.0 | True | 1.0 | 1.0 | fully_conserved |
| TGFB1 | 4 | M3 — Ciliary signaling | -0.0078920584891604 |  | True | 0.0 | False | 0.0 | 0.0 | divergent |
| TGFB3 | 2 | M1 — Structural assembly | 0.0008651026432723 |  | False |  | True | 1.0 | 1.0 | partially_conserved |
| TOM1L2 | 2 | M1 — Structural assembly | 0.134734477506743 |  | False |  | False |  |  | no_cross_species_data |
| TP53BP1 | 2 | M3 — Ciliary signaling | -0.0089294813217911 |  | False |  | False |  |  | no_cross_species_data |
| TPD52 | 3 | M2 — Intraflagellar trafficking | -0.0391281545340961 |  | True | 1.0 | False |  | 1.0 | partially_conserved |
| TPPP3 | 3 | M1 — Structural assembly | 0.0001617980309619 |  | False |  | True | 1.0 | 1.0 | partially_conserved |
| TRAPPC10 | 2 | M3 — Ciliary signaling | 0.0914655506248085 |  | False |  | False |  |  | no_cross_species_data |
| TRAPPC3 | 2 | M3 — Ciliary signaling | -0.0039820044214439 |  | False |  | False |  |  | no_cross_species_data |
| TRPV4 | 2 | M3 — Ciliary signaling | 0.0038677917947566 |  | False |  | True | 1.0 | 1.0 | partially_conserved |
| TSC1 | 2 | M1 — Structural assembly | -0.0019738121766784 |  | False |  | False |  |  | no_cross_species_data |
| TTK | 2 | M1 — Structural assembly | 0.0012230104092306 |  | False |  | False | 1.0 | 1.0 | partially_conserved |
| TUBA1A | 4 | M1 — Structural assembly | 0.0361452532782083 |  | True | 1.0 | False |  | 1.0 | partially_conserved |
| TUBA1B | 4 | M1 — Structural assembly | 0.0356333718989338 |  | False |  | False | 1.0 | 1.0 | partially_conserved |
| TUBA1C | 3 | M1 — Structural assembly | 0.0309955550242861 |  | False |  | False |  |  | no_cross_species_data |
| TUBA4A | 2 | M1 — Structural assembly | 0.0048953122112009 |  | False |  | False |  |  | no_cross_species_data |
| TUBB | 4 | M1 — Structural assembly | -0.0060714745610534 |  | False |  | False |  |  | no_cross_species_data |
| TUBB1 | 2 | M1 — Structural assembly | -0.0029118951906222 |  | False |  | False | 0.0 | 0.0 | divergent |
| TUBB2A | 2 | M1 — Structural assembly | -0.0091908130166308 |  | False |  | False |  |  | no_cross_species_data |
| TUBB2B | 2 | M1 — Structural assembly | -0.0036896156234322 |  | True | 1.0 | False |  | 1.0 | partially_conserved |
| TUBB4B | 5 | M1 — Structural assembly | -0.0058697368897978 |  | True | 0.0 | True | 0.0 | 0.0 | divergent |
| TUBB6 | 4 | M1 — Structural assembly | 0.0929218050359418 |  | True | 1.0 | False |  | 1.0 | partially_conserved |
| TUBGCP5 | 2 | M1 — Structural assembly | -0.0134753532852809 |  | False |  | False |  |  | no_cross_species_data |
| TXNDC15 | 2 | M1 — Structural assembly | -0.0148139586668566 |  | False |  | False |  |  | no_cross_species_data |
| UBE2L3 | 2 | M1 — Structural assembly | 0.0091408230091928 |  | False |  | False |  |  | no_cross_species_data |
| UGGT1 | 2 | M1 — Structural assembly | 0.0220709513623696 |  | False |  | False |  |  | no_cross_species_data |
| USP14 | 2 | M4 — Turnover and maintenance | 0.108921128185419 |  | False |  | False |  |  | no_cross_species_data |
| USP8 | 2 | M4 — Turnover and maintenance | -0.0071636303511916 |  | False |  | False |  |  | no_cross_species_data |
| USP9X | 2 | M4 — Turnover and maintenance | 0.230934827184393 |  | False |  | True | 1.0 | 1.0 | partially_conserved |
| UTRN | 3 | M1 — Structural assembly | 0.416823701670084 | up | False |  | True | 1.0 | 1.0 | partially_conserved |
| VCP | 2 | M3 — Ciliary signaling | 0.291985561031824 | up | False |  | True | 1.0 | 1.0 | partially_conserved |
| VDAC3 | 2 | M3 — Ciliary signaling | 0.027078645826206 |  | False |  | False |  |  | no_cross_species_data |
| VPS35 | 2 | M4 — Turnover and maintenance | -0.0088577052364596 |  | False |  | False | 0.0 | 0.0 | divergent |
| VPS37B | 2 | M2 — Intraflagellar trafficking | 0.0450255592993314 |  | False |  | False |  |  | no_cross_species_data |
| VPS37C | 2 | M1 — Structural assembly | 0.0232938277021712 |  | False |  | False |  |  | no_cross_species_data |
| WASF3 | 2 | M1 — Structural assembly | 0.0257824047802866 |  | False |  | True | 1.0 | 1.0 | partially_conserved |
| WWTR1 | 2 | M1 — Structural assembly | 0.436858037221669 | up | False |  | False |  |  | no_cross_species_data |
| XPO1 | 2 | M2 — Intraflagellar trafficking | -0.0014628658611674 |  | False |  | False | 0.0 | 0.0 | divergent |
| YAP1 | 2 | M1 — Structural assembly | 0.155784226176701 |  | False |  | True | 1.0 | 1.0 | partially_conserved |
| YWHAE | 2 | M1 — Structural assembly | 0.0861541912161417 |  | False |  | True | 1.0 | 1.0 | partially_conserved |
| YWHAG | 2 | M1 — Structural assembly | 0.040275130319115 |  | False |  | False |  |  | no_cross_species_data |
| YWHAQ | 2 | M1 — Structural assembly | 0.0918217818654273 |  | False |  | True | 1.0 | 1.0 | partially_conserved |
| ZFAND2B | 2 | M1 — Structural assembly | -0.0104651336375756 |  | False |  | False |  |  | no_cross_species_data |
| ZNF330 | 2 | M1 — Structural assembly | -0.0071202483530598 |  | False |  | True | 0.0 | 0.0 | divergent |
| ZNF423 | 2 | M1 — Structural assembly | 0.0544881453161115 |  | False |  | False | 1.0 | 1.0 | partially_conserved |
